# Hepatocyte UBE4A protects against MASLD and MASH by maintaining ER homeostasis via UBXN4 interaction

**DOI:** 10.64898/2026.10.07.757434

**Authors:** Naga Lakshmi Challa, Vineet Babu, Molee Chakraborty, Matthew Hayes, Gaurav Kumar, Kyle S. McCommis, Craig W Vander Kooi, Danielle Carpenter, Isabela Ramos, Shivankitha Kattekola, Ameya Padakanti, Shikara Patel, Kaitlyn Mills, Sanchita Gupta, Louise Lantier, Brent A. Neuschwander-Tetri, Anutosh Chakraborty

## Abstract

Molecular understanding of the regulation of endoplasmic reticulum (ER) functions in healthy and MASLD livers is limited. Here, we elucidate that Ube4A, a cytosolic E3 ubiquitin ligase, protects against MASLD development by maintaining ER homeostasis during nutritional stress. Hepatocyte-Ube4A deletion (HUKO) augments Gubra-Amylin-diet (GAN-D)-induced steatosis, inflammation, insulin resistance, ER stress and markers of steatohepatitis of the liver in both sexes of mice. Proteomic studies reveal that in addition to altered metabolic and fibrogenic pathways, levels of many ER proteins are also changed in HUKO livers. Impaired ER functions sensitize HUKO livers to nutrition-or chemical (tunicamycin)-induced ER stress. Moreover, HUKO primary hepatocytes exhibit increased ER stress. Furthermore, Ube4A maintains ER homeostasis by facilitating ER-associated protein degradation (ERAD). The lipogenic protein SCD1 is identified as one of the ERAD targets of Ube4A. SCD1 stability and triglyceride accumulation are amplified in HUKO primary hepatocytes. The liver Ube4A-interactome studies identified the ERAD mediator UBXN4 as a major Ube4A interacting protein. Alphafold3-based molecular modelling and interaction site mapping analyses pinpointed the Ube4A binding residues for UBXN4. ER-localized Ube4A-UBXN4 complex facilitates ERAD (e.g., SCD1) and UBXN4-binding deficient Ube4A mutants fail to enhance this process. Finally, Ube4A and UBXN4 are downregulated in the livers of human MASLD and MASH patients and this complex is diminished in GAN-D-fed mouse livers. In conclusion, Ube4A preserves ER homeostasis by maintaining ERAD via its interaction with UBXN4, which maintains liver health. Enhancing levels and/or preserving interaction of these proteins may offer new therapeutic strategies for MASLD treatment.

## Introduction

MASLD (metabolic dysfunction-associated steatotic liver disease) and steatohepatitis (MASH) are characterized by excess lipid accumulation, inflammation, injury and fibrosis of the liver, which increase the risk of cirrhosis, and hepatocellular carcinoma (HCC)^1–3^. Mechanisms that contribute to the development of MASLD and its progression to MASH have not been fully understood. The endoplasmic reticulum (ER) plays a vital role in directly maintaining hepatic protein and lipid homeostasis. Smooth and rough ER are the sites for lipogenesis and lipogenic protein synthesis, respectively. Moreover, most of the circulatory proteins (except for immunoglobulins) are produced by the hepatocyte rough ER^4,5^. Nutritional, inflammatory and hormonal stress disrupts ER protein and lipid handling in hepatocytes. Acute stress activates the adaptive unfolded protein response (UPR)^6,7^, which enhances refolding of misfolded proteins to restore ER homeostasis^8,9^. However, in chronic nutritional stress, increased free fatty acids (FFAs) enhance ER protein misfolding^10^. Also, FFA-derived triglycerides (TAGs) form more cytosolic lipid droplets (LDs), a defining feature of MASLD^11^. Moreover, chronic increase in the levels of FFAs cause lipotoxicity-induced hepatocyte ER stress and cell death by increased activation of cell death-inducing proteins CHOP and JNK/NFκB^9,12–14^. Studies showing that alleviating ER stress ameliorates MASLD features in mice support this concept^15^.

Studies in yeast and mammalian cell lines have established that ER-associated protein degradation (ERAD) protects cells from ER stress and cell death caused by accumulation of misfolded and aggregated proteins^16,17^. When ER proteins get irreparably misfolded, the ER chaperone BiP presents them to the ERAD complex, which facilitates their retrotranslocation from the ER, ubiquitination and proteasomal degradation in the cytosol^16,17^. UBXN4 (UBX domain containing protein 4) is an ER-anchored protein with a C-terminal transmembrane (TM) domain, keeping both its termini hanging in the cytosol^18^. The N-terminal ubiquitin-regulatory X (UBX) and thioredoxin fold (ThF) domains of UBXN4 mediate protein-protein interactions^19^. UBXN4 interacts with the ERAD-ATPase p97/VCP through its UBX domain^18^, and this complex mediates ERAD in cancer cells^18,20^.

Due to the highly secretory nature of hepatocytes^4,5^, the likelihood of protein misfolding and aggregation-induced ER stress is extremely high in these cells. For example, the lipoprotein ApoB-100 is a well-characterized ERAD substrate in hepatocytes^21^. Moreover, ERAD degrades properly folded lipogenic proteins in hepatocytes, which are constitutively expressed at levels higher than needed to meet the demand^22^. Yet, limited information is available on the mechanisms that regulate ERAD in healthy and MASLD conditions^5,22^. Although the transmembrane ERAD-E3 ligases gp78^23,24^, HRD1^25,26^ and March6^27^ have been partly characterized, the impact of cytosolic E3 ligases in hepatocyte-ERAD, ER stress, and MASLD and MASH development is largely unknown.

A few (i.e., 6-7), mammalian E3 ligases contain a unique U-Box catalytic domain, resembling the common RING domain but lacking the Zn^2+^-coordinating residues^28^. Ube4A is such a U-Box domain containing E3 ligase^28–31^. Impaired function of Ube4A has been implicated in metabolic disease. The *Ube4A* promoter region is hypermethylated in obesity^32,33^, and *Ube4A* has been found to be mutated in the coding region in some obese autistic and obese type-2 diabetic patients^34,35^. For example, loss-of-stability mutations in *Ube4A* were observed in an obese-intellectually disabled population^36^. Ube4A is usually highly expressed in the liver^37^ but has been shown to be substantially downregulated in MASLD and MASH patients^38^. These studies strongly suggest that Ube4A may play a protective role in obesity and MASLD. Indeed, we have shown that whole-body Ube4A deletion (UKO) exacerbates high fat diet (HFD)-induced obesity, insulin resistance and hepatic steatosis in mice^37^. However, to what extent hepatocyte-Ube4A maintains liver and systemic metabolic homeostasis and protects the liver from lipotoxic injury is unknown. Moreover, the mechanisms by which this novel E3 ligase regulates lipid metabolism have also not been determined. The mechanistic studies described in this manuscript aim to address these important gaps in knowledge.

## Results

The pathological accumulation of LDs in hepatocytes is a key early feature of MASLD^39^. To determine whether hepatocyte-specific Ube4A deletion (HUKO) exacerbates hepatic steatosis and promotes the development of steatohepatitis in vivo, the HUKO mouse model was developed (Figure 1A) and the response to nutritional stress was evaluated in control LoxP and HUKO mice. Male mice were fed the Gubra-Amylin diet (GAN-D) for 8 and 16 weeks to assess the early development of hepatic steatosis and steatohepatitis.

**Figure 1:**
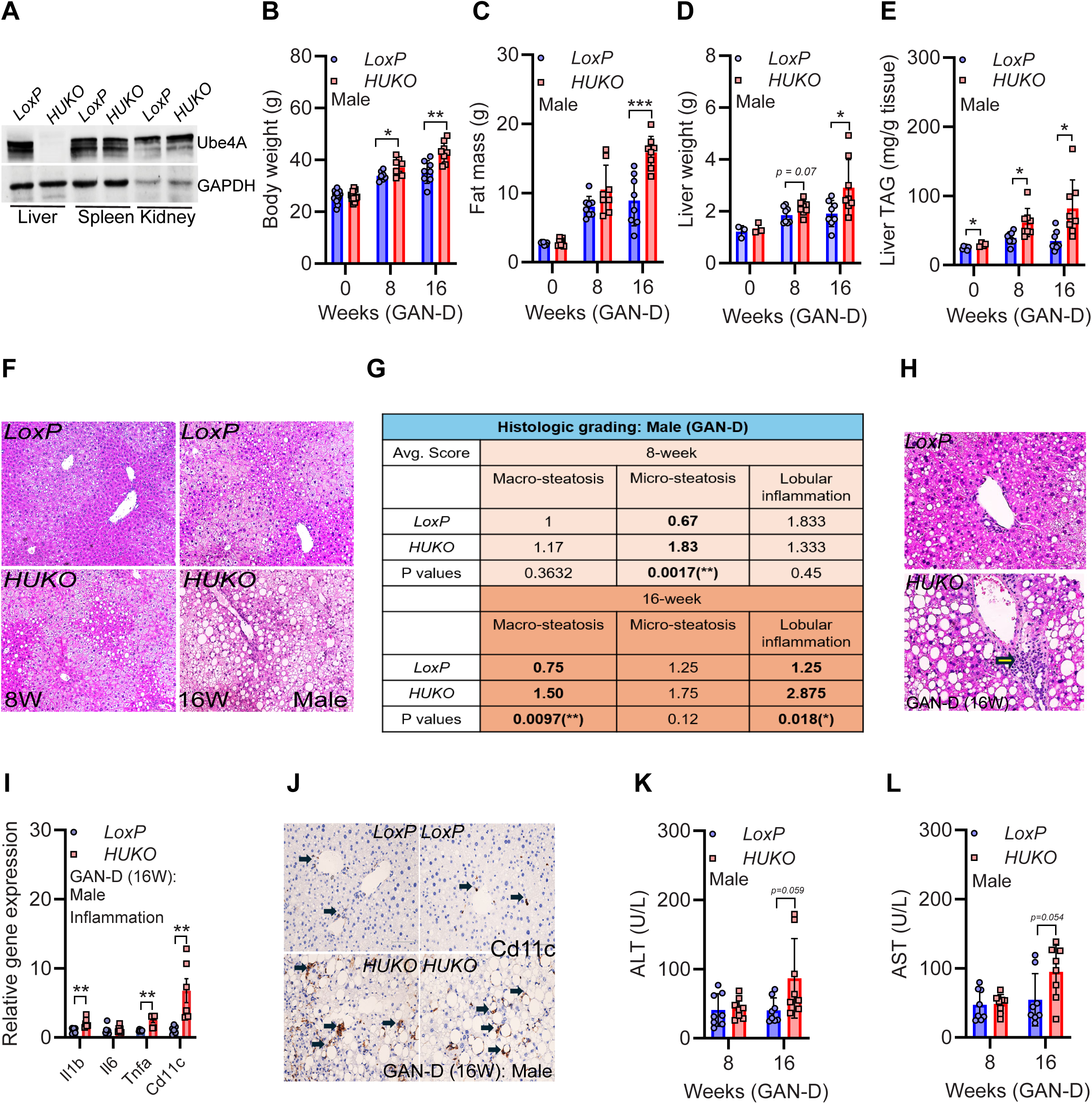
Hepatocyte-specific Ube4A deletion (HUKO) augments GAN diet-induced steatosis, inflammation and injury of the liver in male mice. A. Representative immunoblot analysis confirming the deletion of Ube4A in liver tissue from liver-specific Ube4A knockout mice maintained on a chow diet. B-C. Change in body weight and fat mass in male mice fed the GAN-D for 8 and 16 weeks (N = 8/cohort). D-E. GAN-D–fed male mice exhibited increased liver weight and hepatic TAG levels. F. Representative images of H&E-stained liver sections from male LoxP and HUKO mice following GAN-D feeding. Magnification - 10X, scale bar - 100 µm. G. Histologic grading of macrovesicular steatosis (Macro), microvesicular steatosis (Micro) and lobular inflammation in GAN-D-fed LoxP and HUKO mice (N= 8 per cohort) and are shown with their corresponding p-values. H. Accumulation of inflammatory immune cells (indicated by arrows) in livers of 16-week-GAN-D-fed LoxP and HUKO mice. Magnification - 20X, scale bar - 100 µm. I. Relative hepatic mRNA expression of inflammatory markers in 16-week-GAN-D-fed LoxP and HUKO livers (N = 6/cohort). J. Representative IHC of CD11c in liver tissue from mice fed the GAN-D diet for 16 weeks. CD11c-positive areas are indicated by arrows. Magnification - 20X, scale bar – 50 µm. K-L. Serum levels of AST and ALT in LoxP and HUKO mice (N = 8/cohort).

GAN-D feeding increased body and fat mass in HUKO mice to a higher extent than LoxP mice (Figures 1B-1C). Liver-weight, hepatic TAG content and LDs were also increased in HUKO mice (Figures 1D-1F). A semiquantitative histologic scoring system^40^ showed significantly increased micro-steatosis in 8-week-GAN-D-fed HUKO livers, progressing to macro-steatosis after 16 weeks (Figures 1F-1G). Moreover, serum TAG levels were increased in GAN-D-fed HUKO mice (Figures S1A). LoxP and HUKO male mice exhibited similar oxygen consumption (VO_2_), energy expenditure (EE), energy preference (respiratory exchange ratio – RER) and activity levels (Figures S1B-S1F) suggesting increased hepatic steatosis in HUKO mice is not due to altered whole body energy metabolism but is a consequence of hepatic metabolic dysfunction.

Livers from 16-week-GAN-D-fed HUKO male mice exhibited increased inflammation (Figures 1G-1H). Accordingly, hepatic expression of the inflammatory markers *Cd11c*, *Il1b*, and *Tnfα* were significantly elevated in HUKO livers in this condition (Figures 1I-1J). In addition, HUKO mice displayed amplified liver injury compared to LoxP control GAN-D-fed mice, evidenced by increased serum levels of alanine and aspartate aminotransferase enzymes (ALT and AST) (Figures 1K-1L). These results demonstrate that hepatic Ube4A deficiency not only aggravates steatosis, but also markedly increases the markers of steatohepatitis in the setting of nutritional stress.

Fibrogenic pathway markers and the development of measurable fibrosis were also assessed in 16- and 24-week GAN-D-fed HUKO male mice. Despite increased expression of fibrogenic genes (Figures S1G-S1H), collagen deposition was only marginally higher in HUKO livers (Figures S1I-S1J). Interestingly, body weight, fat mass, liver-weight and TAG content became similar in LoxP and HUKO GAN-D-fed male mice at 24 weeks (Figure S1K-S1N). Serum TAG levels were still higher in HUKO mice, indicating increased lipid-export to combat hepatic lipotoxicity in these mice (Figure S1O). In summary, hepatocyte-specific Ube4A deletion augments GAN-D-induced steatosis and early induction of inflammation and injury of the liver in male mice.

In humans, MASLD occurs about equally in males and females^41^, but in mouse models it is typically more prominent in males compared to females. Therefore, we separately assessed the impact of hepatocyte-Ube4A deletion on female mice, which take a slightly longer duration of nutritional stress to develop obesity and MASLD^37,42–45^. After 32 weeks of GAN-D feeding, both LoxP and HUKO female mice exhibited similar body weight, fat mass and lean mass (Figures S2A-S2C). Yet female HUKO livers showed increased weight and TAG accumulation (Figures 2A-2C). Serum levels of TAG were also higher in female HUKO mice (Figure 2D). Female HUKO livers exhibited features of steatohepatitis with increased expression of inflammatory markers (Figure 2E). Most importantly, female HUKO mice developed more fibrosis than GAN-D-fed LoxP controls (Figures 2F-2G). Accordingly, fibrotic marker genes and proteins including the activated hepatic stellate cell (HSC) marker αSMA (alpha smooth muscle actin, gene: *Acta2*) were upregulated in female HUKO livers (Figures 2H-2K). Consistent with these findings, serum levels of the liver injury markers ALT and AST were increased in female HUKO mice (Figures 2L-2M). Thus, GAN-D-induced MASLD and MASH features are more severe in female HUKO mice than wild-type GAN-D fed mice.

**Figure 2:**
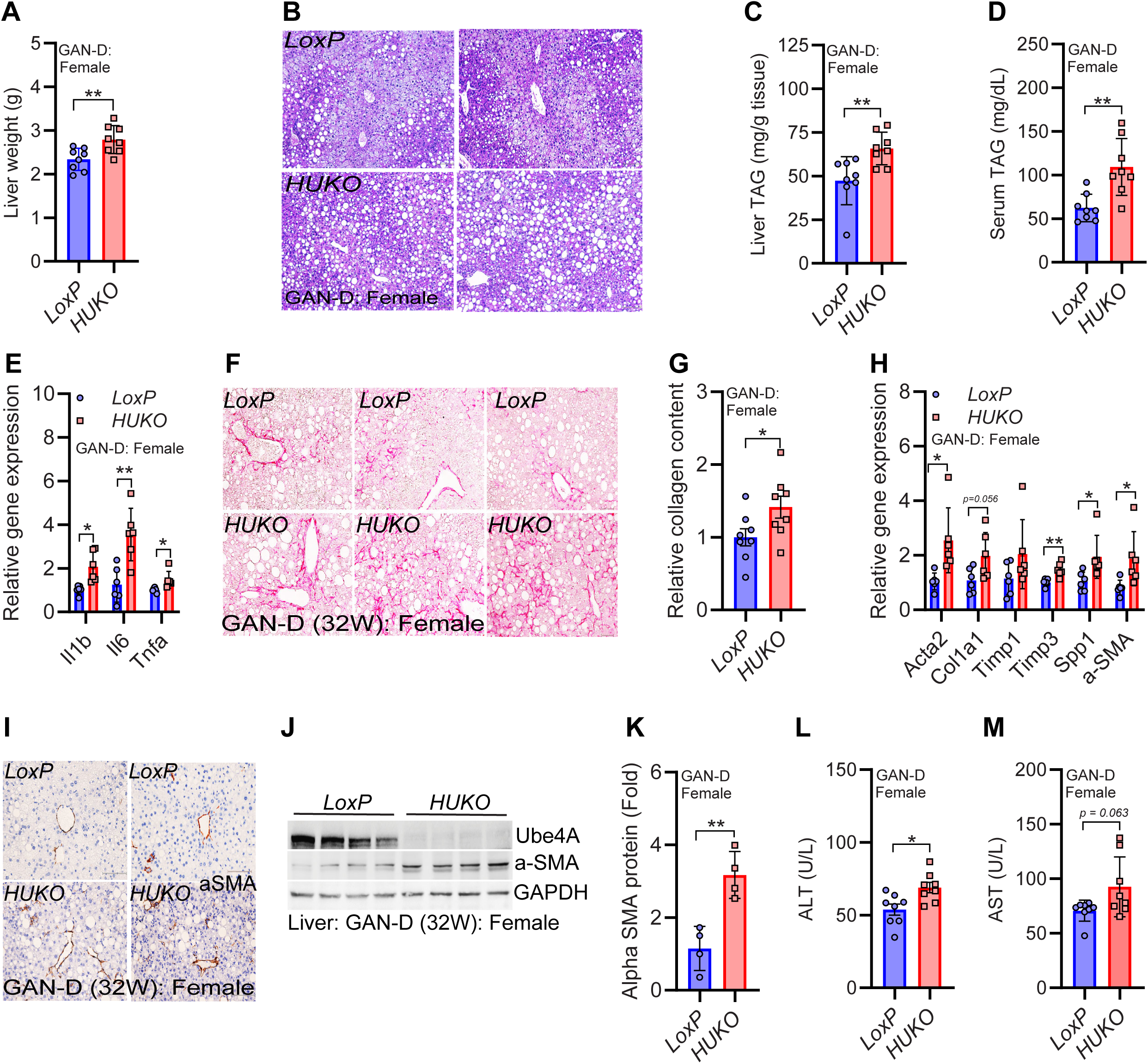
GAN-D-induced MASLD features are amplified in female HUKO mice. A. Liver weight in 32-week-GAN-D-fed female mice (N=8/cohort). B. Representative images of H&E-stained liver sections from female LoxP and HUKO mice. Magnification - 10X, scale bar - 100 µm. C. Liver TAG content in female LoxP and HUKO mice (N = 8/cohort). D. Serum TAG levels in female LoxP and HUKO mice. E. Relative expression of inflammatory marker genes in female LoxP and HUKO mice (N=6/cohort). F-G. Representative Picrosirius-red staining of liver sections along with the quantification in female LoxP and HUKO mice. Magnification - 10X, scale bar - 100 µm. H. Hepatic expression levels of fibrotic gene markers in female LoxP and HUKO mice (N= 6/cohort). I. IHC for α-SMA in liver sections from female LoxP and HUKO mice. Magnification - 20X, scale bar - 50 µm. J-K. Immunoblots analysis and quantification for α-SMA in female LoxP and HUKO livers. L-M. Serum levels of AST and ALT in female LoxP and HUKO mice (N = 8 per cohort).

HFD feeding impairs disposal of exogenous glucose and insulin-induced glucose uptake (GTT and ITT) in male and female whole-body Ube4A-KO (UKO) mice^37^. To test whether hepatocyte-Ube4A deletion also worsens insulin resistance, hyperinsulinemic-euglycemic clamp experiment^46–48^ was performed on 16-week-GAN-D-fed male mice. Blood glucose, glucose infusion rate (GIR) and plasma insulin levels were similar in both genotypes (Figures 3A and S3A-S3B). The rate of endogenous glucose production during clamp (endoRa) was slightly lower in HUKO mice whereas glucose disappearance (Rd) was similar in both genotypes (Figures 3B-3C). 2-deoxyglucose (2-DG) uptake (Rg) was also similar in most metabolic tissues except the brown adipose tissue (BAT), which exhibited significantly impaired glucose uptake in HUKO mice (Figures 3D and S3C) suggesting an impact of liver metabolism on BAT. GTT and ITT, performed on 12-13-week-GAN-D-fed male mice, also showed no significant differences between genotypes (Figures S3D-S3E). Thus, hepatocyte-specific Ube4A deletion does not cause early-onset insulin resistance (12-16-week-GAN-D-fed) in male mice except in BAT.

**Figure 3:**
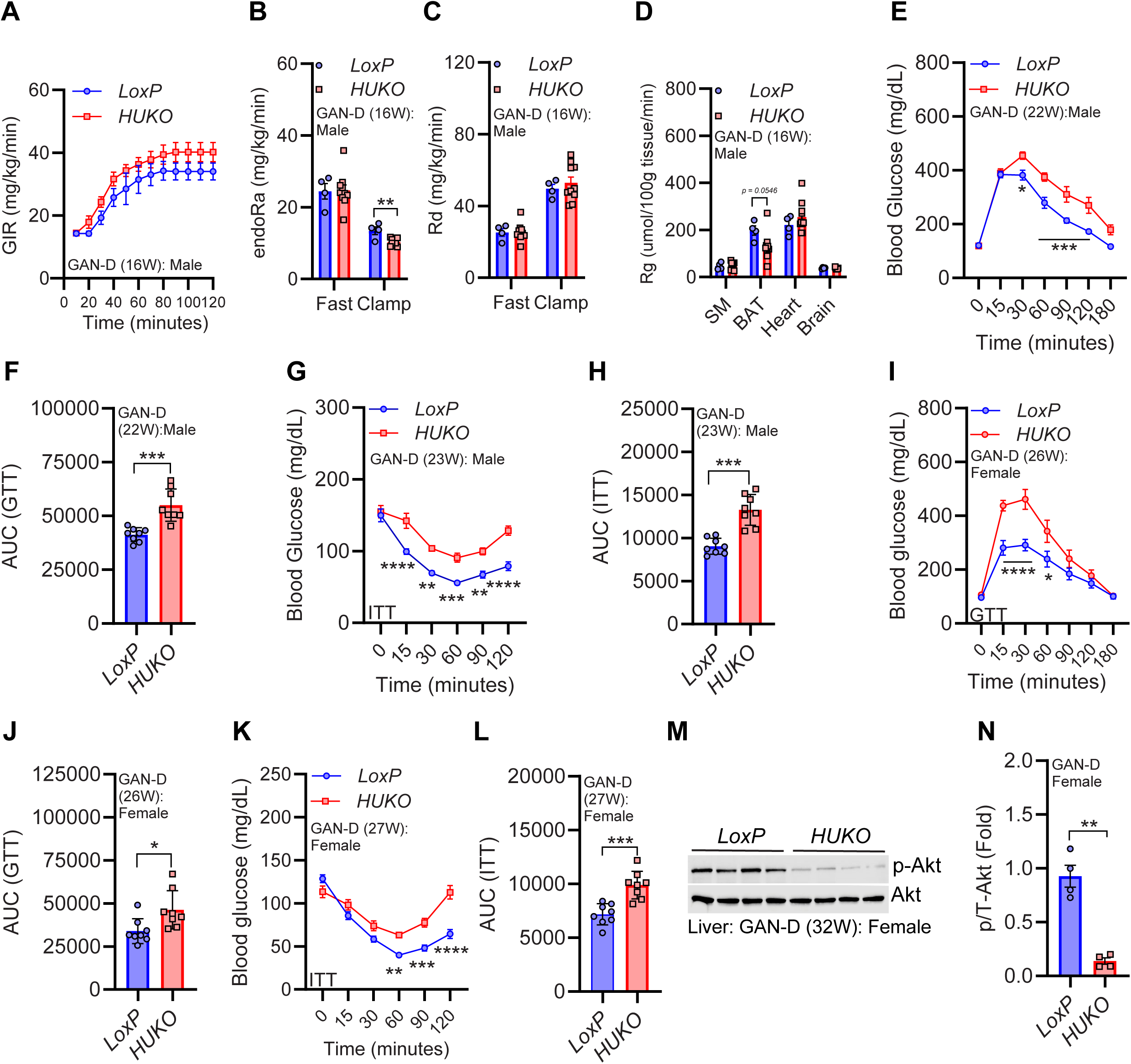
GAN-D-fed HUKO mice exhibit delayed onset insulin resistance. A. Glucose infusion rate (GIR) in 16-week-GAN-D-fed male LoxP and HUKO mice (N = 4 - WT and 10 – HUKO). B-C. Clamp-derived rates of endogenous glucose production (endoRa) and whole-body glucose disappearance (Rd) in LoxP and HUKO male mice. D. Tissue-specific 2-DG uptake (Rg) in metabolic tissues of LoxP and HUKO mice. E-F. Raw and AUC values of GTT in 22-week-GAN-D-fed male LoxP and HUKO mice (n = 8/cohort). G-H. Raw and AUC values of ITT in 23-week-GAN-D-fed male LoxP and HUKO mice (n = 8/cohort). I-J. Raw and AUC values of GTT in 26-week-GAN-D-fed female LoxP and HUKO mice (n = 8/cohort). K-L. Raw and AUC values of ITT in 27-week-GAN-D-fed female LoxP and HUKO mice (n = 8/cohort). M-N. Immunoblots and densitometry of phosphorylated Akt (S473) in 32-week-GAN-D-fed female LoxP and HUKO livers.

However, GTT and ITT, performed on 22-23-week-GAN-D-fed male mice and 26-27-week-GAN-D-fed female mice showed increased insulin resistance in HUKO mice (Figures 3E-3L). Accordingly, stimulatory phosphorylation (S473) of the insulin effector protein kinase Akt was substantially lower in HUKO livers (Figures 3M-3N). Thus, the development of hepatic insulin resistance occurs in HUKO mice in the setting of inflammation/injury response rather than steatosis.

To determine which tissues are responsible for early-onset insulin resistance in HFD-fed whole-body Ube4A-KO (UKO) mice^37^ (as opposed to delayed insulin resistance in GAN-D-fed HUKO mice described above), hyperinsulinemic-euglycemic clamp was performed in HFD-fed UKO mice. Blood glucose levels were higher, and GIR was lower in 12-week-HFD-fed UKO mice despite higher plasma insulin levels than WT (Figures S3F-S3H). EndoRa (glucose production) was higher whereas the rate of glucose disappearance (Rd) was lower in UKO mice (Figures S3I-S3J). Insulin stimulates glucose disappearance in WT (Rd shifted higher with more insulin) but not in UKO mice, indicating severe peripheral insulin resistance in HFD-fed UKO mice (Figure S3K). Like HUKO mice, 2-DG uptake (Rg) was most impaired in the BAT of UKO mice followed by heart and muscle (Figures S3L and S3M). Thus, early onset, severe insulin resistance in UKO mice is due to impaired glucose uptake in BAT, heart, and skeletal muscle, an effect not observed in HUKO mice, which showed delayed onset insulin resistance.

The most striking and early observed phenotype of HUKO mice is hepatic steatosis. Therefore, we further characterized this feature. Untargeted lipidomic studies in 16-week-GAN-D-fed male mice were performed to detect alterations in specific lipid species in HUKO livers (Figures 4A-4B, S4A-4D). As expected, the levels of glycerolipids (e.g., TAG, DAG) were increased (total glycerolipids, HUKO vs LoxP: 64% vs 46%, TAG: 42% vs 31% and DAG: 22% vs 15%) in GAN-D-fed HUKO livers compared to GAN-D-fed LoxP controls (Figures 4A-4B and S4A-S4B). However, the relative abundance of glycerophospholipids (e.g., membrane lipids such as phosphatidylcholine [PC] and phosphatidylethanolamine [PE]) were decreased in GAN-D-fed HUKO livers compared to LoxP controls (total glycerophospholipids, HUKO vs LoxP: 33% vs 51%; PC: 29% vs 43% and PE: 4% vs 7%) (Figures 4B and S4B). The most increased (∼11-fold) TAG species in HUKO livers was tripalmitin (16:0_16:0_16:0), a major component of GAN-D, and marker of MASLD, de novo lipogenesis and activity of fatty acid synthase (FAS)^49^ (Figure S4E). Moreover, several TAG (e.g., 14:1_16:1_18:1) and DAG (e.g., 16:0_18:1) species with monounsaturated fatty acids (MUFA) were also increased in the HUKO livers (Figure S4E).

**Figure 4:**
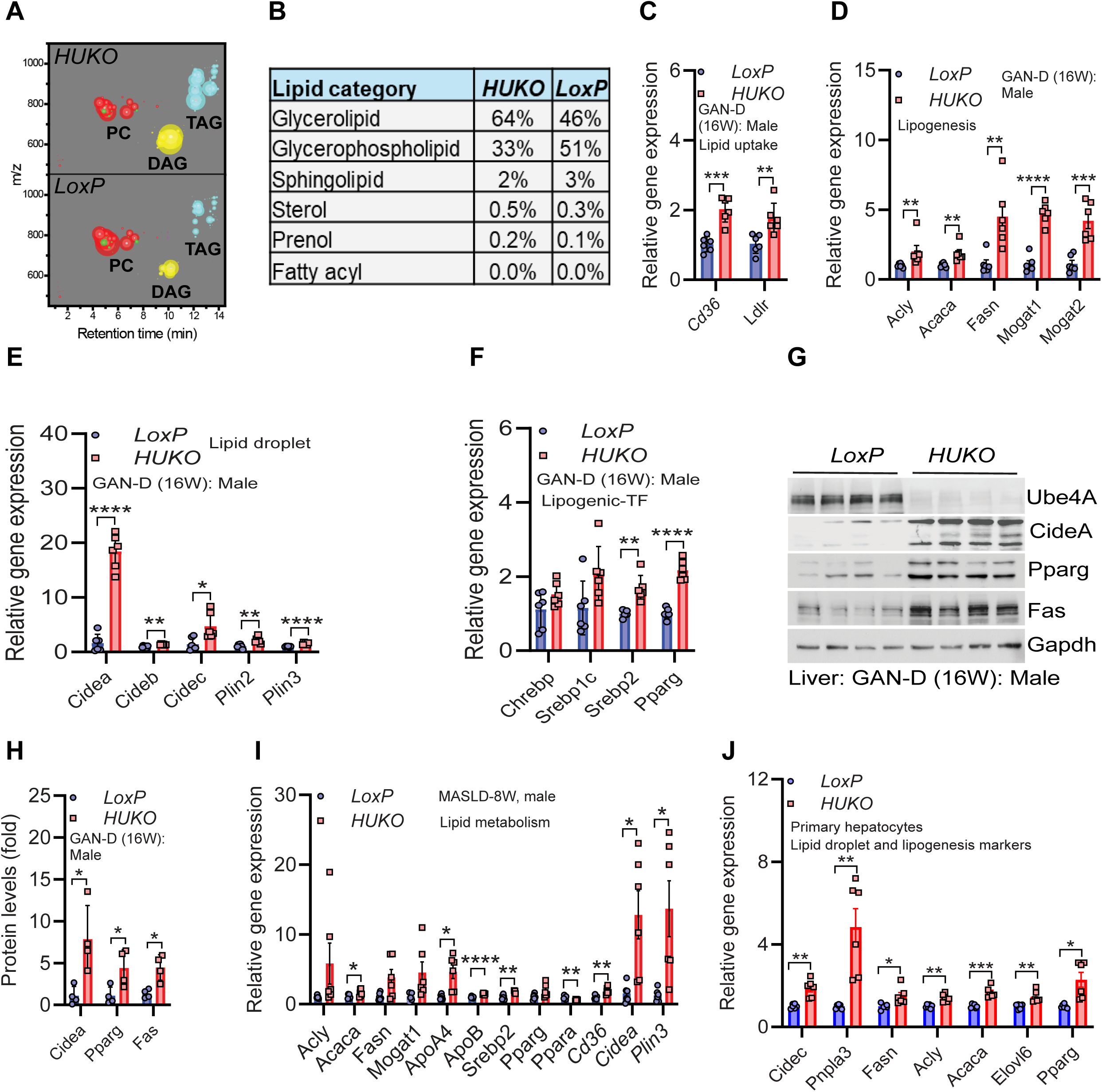
Hepatocyte-Ube4A deletion upregulates lipid accumulation pathways, leading to increased TAG accumulation and hepatic steatosis in GAN-D-fed mice. A-B. Lipid class distribution in 16-week-GAN-D-fed male LoxP and HUKO liver samples by mass spectrometry (N = 5 per cohort). C-F. Relative hepatic expression of genes involved in, lipogenesis, LD formation, and lipogenic transcription factors in 16-week-GAN-D-fed male LoxP and HUKO mice (N=6 per cohort). G. Immunoblot analysis of lipogenic markers in the livers of male LoxP and HUKO mice. H. Relative hepatic expression of lipid metabolic genes in 8-week-GAN-D-fed male LoxP and HUKO mice (N = 6 per cohort). I. Relative expression of lipid metabolic genes in LoxP and HUKO primary hepatocytes (N = 6 experimental replicates from 2 mice per group). J. Hepatocyte TAG levels LoxP and HUKO primary hepatocytes (N = 5 experimental replicates from 2 mice per group).

Increased hepatic steatosis was also supported by gene and protein expression analyses. Expression of fatty acid biosynthesis, lipogenesis, lipid uptake and LD-stabilizing pathways and their transcription factors were upregulated in 16-week-GAN-D-fed HUKO livers (Figures 4C-4G). VLDL lipoprotein genes such as *ApoB* and *ApoC2*, VLDL assembly protein *Mttp*, and VLDL-export enhancing protein *ApoA4* were also increased in GAN-D-fed HUKO mice, suggesting that TAG export in VLDL is increased to reduce the extent of hepatic steatosis in HUKO mice (Figure S4F). Lipid metabolic genes exhibited similarly higher trends in 8-week-GAN-D-fed HUKO mice, consistent with early hepatic steatosis in HUKO mice (Figure 4H).

HUKO primary hepatocytes exhibited increased expression of the lipogenic markers and greater intracellular TAG accumulation under basal conditions, indicating cell-autonomous role of Ube4A in reducing de novo lipogenesis and hepatocellular lipid storage independent of external influences (Figure 4I-4J).

Although the physiological functions of Ube4A have recently been appreciated^37^, specific information is not available on which proteins/pathways in the liver are regulated by this novel E3 ligase. To identify changes in the abundance of specific liver proteins in the absence of hepatocyte-Ube4A, nano-liquid chromatography tandem mass spectrometry (Nano-LC/MS/MS)^50^ was performed in 16-week-GAN-D-fed liver samples from male mice. This approach identified 441 proteins (∼11.6% of 3810 proteins detected) which were significantly altered in HUKO livers. Pathway enrichment analysis was conducted using significantly altered proteins in HUKO livers. Dot plots summarizing the top 20 significantly enriched KEGG pathways are shown in Figures 5A-5B and Gene Ontology Cellular Component (GO:CC) analysis identifying subcellular localization of the proteins are shown in Figures 5C-5D. A Volcano plot, summarizing the altered protein levels and showing proteins that may explain the HUKO phenotype are presented in Figure S5A.

**Figure 5:**
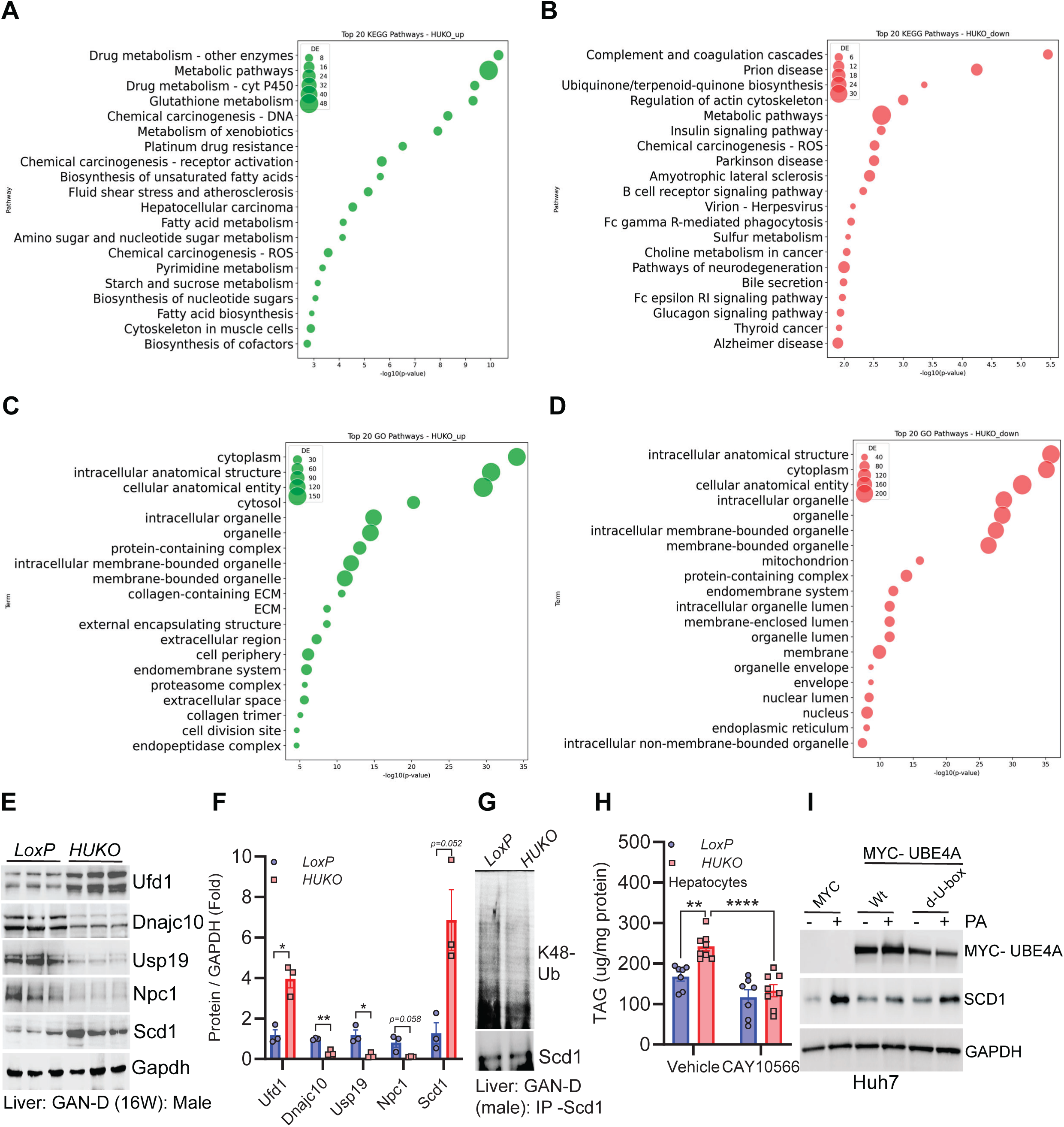
The liver proteome of GAN-D-fed HUKO mice exhibits alterations in MASLD, ER processing and ERAD pathway proteins. A-B. Dot plots summarizing top 20 significantly enriched KEGG pathways in 16-week-GAN-D-fed male HUKO mice (N=4/genotype). C-D. Gene Ontology Cellular Component (GO:CC) analysis for subcellular localization of the proteins identified by proteomics. E-F. Immunoblotting and densitometric quantification of Ufd1, Dnajc10, Usp19, Npc1 and Scd1 in LoxP and HUKO livers. G. Representative immunoblot showing K48-ubiquitination (K48-Ub) levels of Scd1, immunoprecipitated from livers of 16-week-GAN-D-fed male LoxP and HUKO mice. H. TAG levels in LoxP and HUKO hepatocytes under basal conditions with vehicle and SCD1 inhibitor CAY10566 (N = 6-8 experimental replicates from N = 2 mice / genotype). I. Immunoblot analysis of PA-induced SCD1 levels in Myc (control), wt-Ube4A and d-u-box-Ube4A overexpressed Huh7 cells.

KEGG pathway analysis identified several pathways that are crucial for liver health. Among these, upregulation of fatty acid biosynthesis, and downregulation of ubiquinone biosynthesis (related to mitochondrial oxidative phosphorylation) and insulin signaling were most relevant to the increased lipotoxic injury in HUKO livers (Figures 5A-5B). Overall, 55 (∼12.5%) and 13 (∼3%) of the 441 significantly altered proteins are known regulators of lipid metabolism, mitochondrial function and insulin signaling, although it is to be noted that substantial overlap exists among metabolic pathways.

A few key upregulated proteins were lipogenic ChREBP, SCD1 (stearoyl-CoA desaturase-1), ACSL3 (acyl-CoA synthetase long chain family member 3), FAS and PLIN2, and downregulated proteins were cholesterol export proteins NPC1 (Niemann-Pick C1), ABCA1 (ATP binding cassette transporter A1) and oxidative phosphorylation proteins COX5b, NDUFA12 and COQ5 (Figure S5B). Insulin signaling proteins (such as MAPK1, APPL1, AKT2 etc.) were generally downregulated (Figure S5C). Pro-fibrogenic pathways were also upregulated and anti-fibrotic pathways downregulated (except for fibrogenic integrin beta-3, which is downregulated) at the protein level in HUKO mice, consistent with the mRNA data (Figure S5D).

Gene Ontology Cellular Component (GO:CC) analyses show that proteins in various cellular compartments were altered in GAN-D-fed HUKO livers (Figures 5C-5D). A substantial portion of these altered proteins (179 of 441, ∼40.6%) have direct connection with functions of the ER. For example, proteins involved in ER protein folding/processing such as DNAJC10 and CRELD2 and the ER-anchored ubiquitin-specific peptidase USP19 were downregulated in HUKO livers (Figure S5E). Conversely, ERAD mediating proteins such as UFD1, ERLIN-2, BAG6 and UBE2b were upregulated in HUKO livers (Figure S5E). Alterations in the abundance of several proteins (UFD1, DNAJC10, USP19 and NPC1) were validated by immunoblotting (Figures 5E-5F).

An important protein that mechanistically links Ube4A with lipogenesis and ERAD is SCD1. Mono-unsaturation of fully saturated fatty acids such as palmitate (PA) (SFA to MUFA) by the ER-anchored enzyme SCD1 enhances synthesis of TAG (LDs), phospholipids, and cholesteryl esters^51^. In hepatocytes, properly folded and active SCD1 is degraded by ERAD^52^, although the E3 ligase that mediate SCD1 ERAD in the setting of MASLD has not been identified. Protein levels of SCD1 were increased whereas its ubiquitination levels decreased in GAN-D-fed HUKO livers (Figures 5E-5G). Accordingly, HUKO primary hepatocytes, in serum and FA-free media, exhibited increased TAG accumulation under basal conditions (Figure 5H). Moreover, treatment with a selective SCD1 inhibitor CAY10566 reduced TAG in HUKO hepatocytes to the levels of WT hepatocytes, confirming that increased TAG in absence of Ube4A is due to enhanced SCD1-mediated TAG formation (Figure 5H).

Ube4A reduces SCD1 protein levels in a cell-autonomous manner. PA, a known inducer of SCD1 protein^53^, increased SCD1 in GST-overexpressed (control) but not in wt-Ube4A-overexpressed Huh7 cells (Figure 5I). Catalytically inactive (u-box deleted or d-U-box) Ube4A did not reduce PA-induced SCD1 protein level (Figure 5I), suggesting that the catalytic activity of this ligase is essential in reducing SCD1 protein level. Overall, these findings strongly suggest that Ube4A maintains hepatocyte ER protein and lipid homeostasis by enhancing ERAD of the lipogenic protein, which is crucial to control hepatic steatosis.

The above findings suggest that hepatocyte-specific Ube4A deletion disrupts ER protein and lipid homeostasis due to impaired ERAD. Impaired ERAD causes ER stress, which is a key causative factor of MASLD^9,12–14,54^. Therefore, we determined if Ube4A deletion enhances ER stress. Indeed, gene expression of the ER stress-induced cell death protein Chop was progressively increased in GAN-D-fed HUKO mouse livers (Figure 6A). Protein levels of Chop and stimulatory phosphorylation status of the ER stress-induced inflammatory and cell death inducing protein Jnk1 were also increased in both sexes of HUKO mouse livers (Figures 6B-6G). Gene expression of the ER chaperone Bip and the upstream UPR/ER stress pathway genes were similar in both genotypes (Figure S6). Besides nutritional stress caused by the GAN-D feeding, hepatocyte-specific Ube4A deletion also augmented tunicamycin (Tm)-induced ER stress in the livers of chow-fed HUKO mice (Figures 6H-6J). Ube4A deletion inherently triggers ER stress in a cell-autonomous manner as its deletion increased expression of *Chop* along with *Bip* and *Xbp1* in primary hepatocytes under basal conditions (Figure 6K). Likewise, Ube4A knockdown (KD) in Huh7 cells also resulted in increased CHOP, altogether suggesting that loss of Ube4A directly triggers ER stress (Figures 6L-6M).

**Figure 6:**
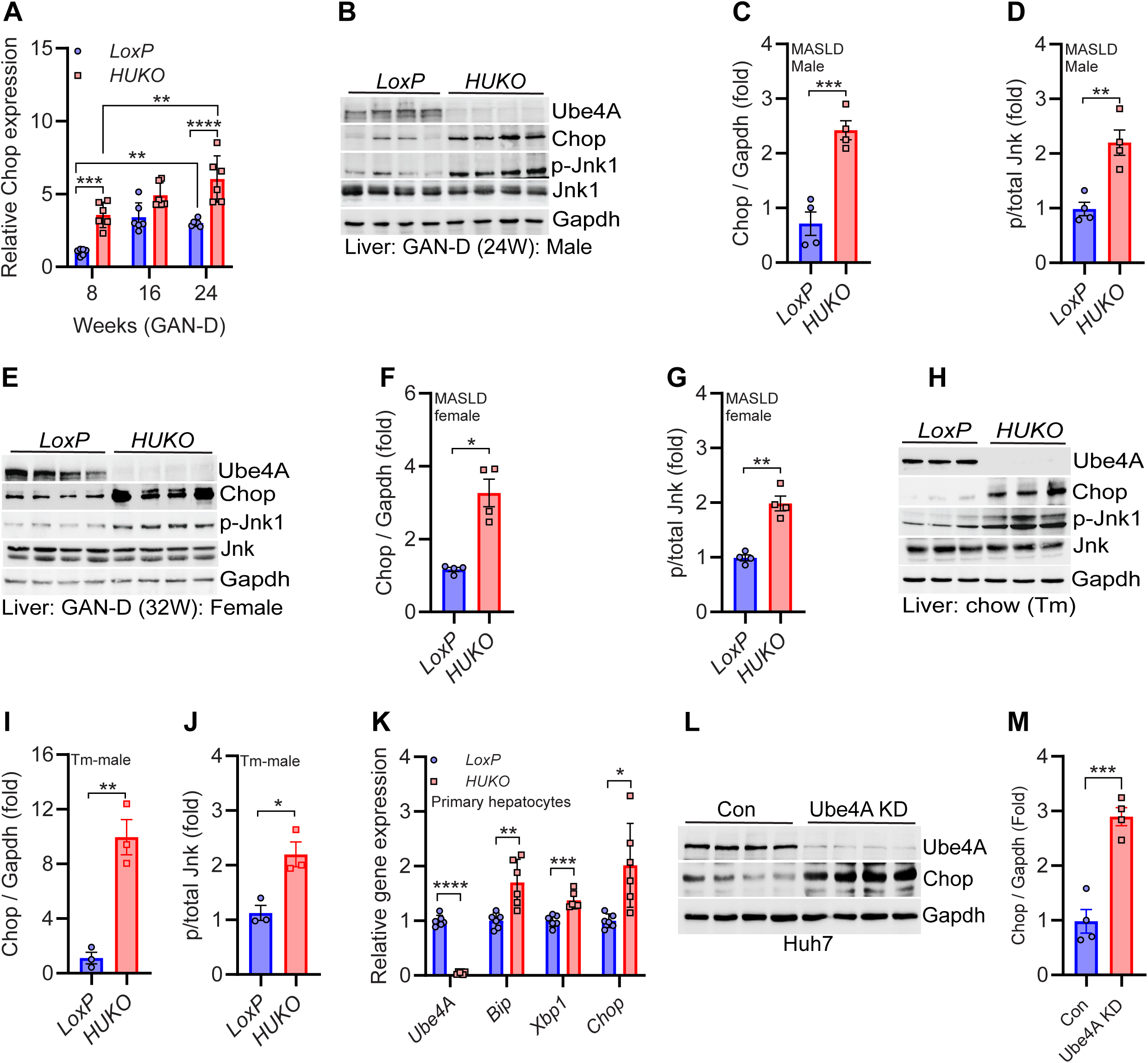
Hepatocyte ER stress is increased in absence of Ube4A. A. Relative mRNA levels of Chop in GAN-D-fed male LoxP and HUKO livers (N=6/cohort). B-D. Immunoblot analysis and quantification for Chop and phosphorylated JNK (p-JNK) in 24-week-GAN-D-fed male LoxP and HUKO livers. E-G. Immunoblot analysis and quantification for Chop and p-JNK in 32-week-GAN-D-fed female LoxP and HUKO livers. H-J. Immunoblot analysis and quantification of Chop and p-JNK in livers from chow-fed LoxP and HUKO male mice treated with vehicle or tunicamycin. K. Relative expression of ER stress genes in LoxP and HUKO primary hepatocytes under basal condition (N = 6 experimental replicates from 2 mice per group). L-M. Immunoblot and quantification of Chop in UBE4A-knockdown (KD) Huh7 cells.

Ube4A is a cytosolic E3 ligase, therefore it can mediate ERAD if it interacts with an ER-anchored, ERAD mediator protein. Therefore, subsequent experiments were designed to identify proteins that might interact with Ube4A to facilitate its role in regulating ER functions. The Ube4A-interactome in chow-fed liver and HEK293 cells, identified by the immunoprecipitation-LC/MS/MS technique, detected UBXN4 as the top interactor of this ligase (Figures 7A, S7A-S7D). Endogenous Ube4A and UBXN4 co-localized in Huh-7 cells (Figure 7B). These proteins also interacted in GAN-D-fed liver samples albeit to a lesser extent likely due to reduced levels of UBXN4 in these conditions (Figures 7C).

**Figure 7:**
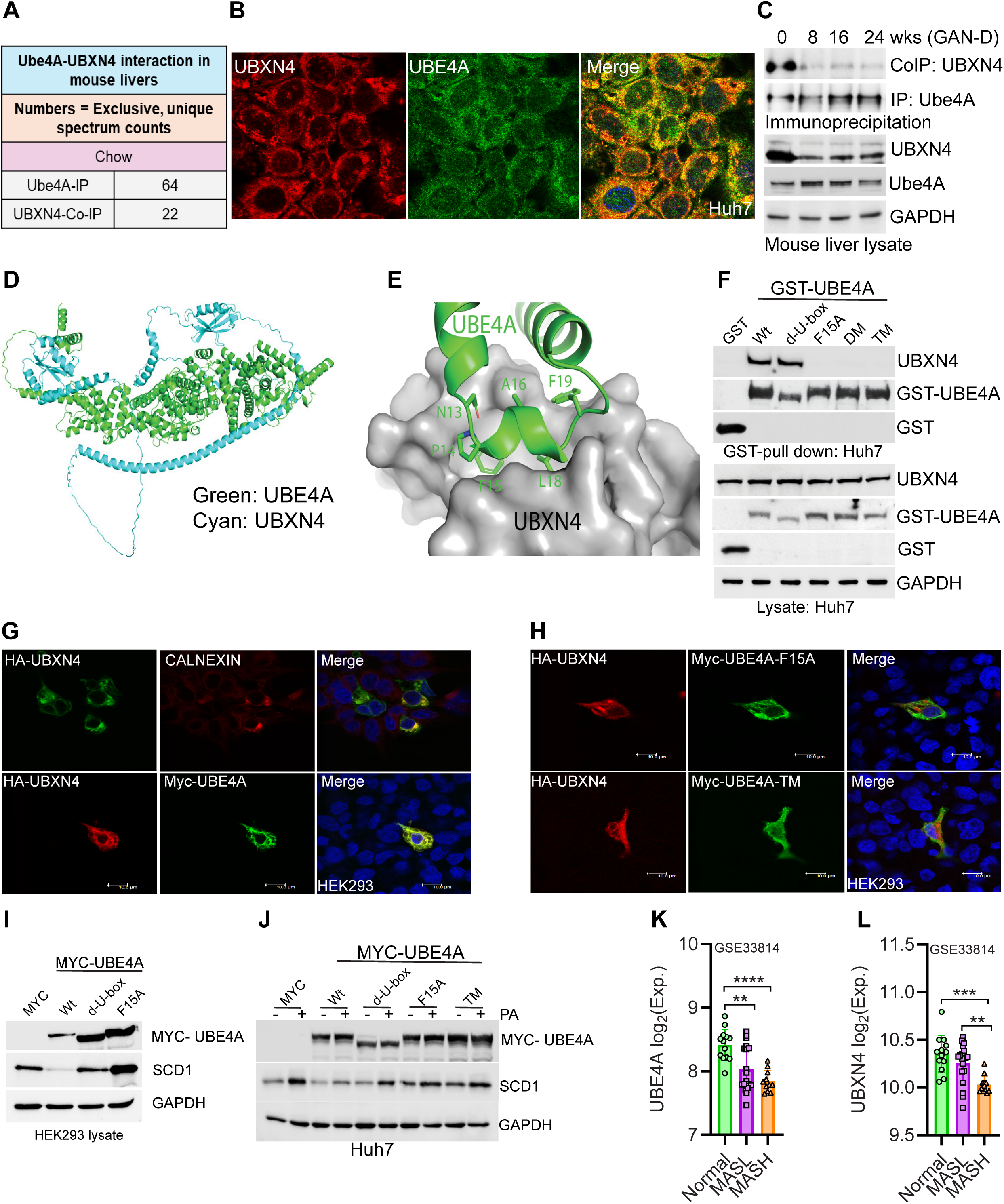
Interaction with the ERAD mediator UBXN4 facilitates ER-localization of Ube4A and degradation of Scd1 in the liver. A. Identification of endogenous UBXN4 and Ube4A in chow-fed mouse liver by Ube4A-immunoprecipitation followed by LC-MS/MS. Residues with post-translational modifications such as oxidation, acetylation etc. are highlighted green. B. Representative confocal images showing co-localization of endogenous UBXN4 and UBE4A in Huh-7 cells. C. Immunoblot analysis showing expression of Ube4A and UBXN4, and their interaction in chow- and GAN-D-fed (8, 16 and 24 weeks) WT mice. D-E. Alphafold3 analysis showing the UBE4A-UBXN4 interaction and the contacts residues of Ube4A. F. GST-pulldown assay showing interaction of endogenous UBXN4 with overexpressed wt and catalytically dead (U-box-deleted or d-U-box) UBE4A in Huh-7 cells. UBXN4 binding efficient (F15M, double mutant - DM and triple mutant – TM) UBE4A mutants failed to bind with UBXN4. G. Top panel. Representative confocal images showing ER-localization of HA-UBXN4 in Huh-7 cells. Calnexin was used as an ER marker. Bottom panel – co-localization of HA-UBXN4 with Myc-UBE4A. Magnification – 100X, scale bar - 10 µm. H. Confocal images show no co-localization of HA-UBXN4 with UBXN4 binding deficient UBE4A mutants F15A and TM, which are dispersed in the cytosol in Huh-7 cells. Magnification – 100X, scale bar - 10 µm. I. Immunoblot analysis of SCD1 protein levels in HEK293 cells overexpressing wt-, d-U-box- or F15A-UBE4A. J. Immunoblot analysis of PA-induced SCD1 expression in Huh7 cells expressing wt-, d-U-box-, F15A- or TM-UBE4A. K-L. UBE4A and UBXN4 expression in human liver transcriptomic datasets across healthy, MASL, and MASH groups.

Studies were then undertaken to identify the specific sites on Ube4A that are crucial for UBXN4 binding-mediated ER localization of this ligase. Alphafold3 predicted a very strong and specific interaction between the N-termini of human Ube4A (residues 1-47, disordered region) and human UBXN4 (residues 1-125, thioredoxin-like ThF domain) (Figures 7D and S7E). Further molecular analysis identified the specific interaction as a high confidence prediction (pTM=0.74) and hydrophobic interactions between F15, L18, F19 of human Ube4A and I12 and I71 of human UBXN4 as the core interface of this complex (Figure 7E). These proposed contact residues are conserved in mice and humans. Deletion and point mutation analyses confirmed that F15A, F15A/L18A (double mutant or DM), and F15A/L18A/F19A (triple mutant or TM) mutants of Ube4A all failed to robustly bind endogenous UBXN4 in Huh7 cells (Figure 7F). Similar observations were made in HEK293 cells (Figure S7F).

Next, we determined the cellular compartment where Ube4A interacts with UBXN4. ER-localization of UBXN4 was evident in HEK293 cells as it co-localized with the ER-membrane protein calnexin (Figure 7G – top panel). Moreover, complete co-localization of overexpressed wt-Ube4A and wt-UBXN4 was observed (Figure 7G – lower panel). However, UBXN4 binding deficient Ube4A mutants failed to co-localize with UBXN4 and thus were dispersed in the cytosol (Figures 7H). These findings indicate that the Ube4A-UBXN4 complex forms at the cytosolic face of the ER.

We further examined whether the Ube4A-UBXN4 complex mediates ERAD. Based on the in vivo results (Figures 5E-5H), we selected SCD1 as a putative ERAD substrate for this complex. Overexpression of wt-Ube4A substantially reduced SCD1 protein level in HEK293 cells whereas catalytically inactive (delta-Ubox or d-Ubox) Ube4A did not exhibit this effect, confirming the essentiality of the E3 ligase activity of Ube4A in SCD1-ERAD (Figure 7I). UBXN4 binding deficient Ube4A (F15A) mutant also failed to reduce SCD1 protein levels, showing the importance of the Ube4A-UBXN4 complex in this process (Figure 7I). Moreover, basal and PA-induced increases in SCD1 were abrogated in wt-Ube4A but not d-Ubox or UBXN4 binding deficient versions of Ube4A in Huh7 cells (Figure 7J).

Finally, to assess the correlation between UBE4A and UBXN4 expressional changes in our mouse models with human MASLD, we analyzed publicly available human liver RNA microarray datasets from control (N = 13) MASL (N = 19) and MASH (N = 12) subjects were studied (GSE33814)^55^. This analysis showed that UBE4A and UBXN4 are downregulated in MASL and MASH patients (Figure 7K-7L).

In summary, we discovered that Ube4A resides at the cytosolic face of the ER by interacting with the ERAD mediator UBXN4. Nutrient-overload enhances misfolding of ER-processed proteins and SCD1 activity, leading to ER stress and hepatic steatosis. The Ube4A-UBXN4 complex protects hepatocytes from these aberrations by enhancing ERAD of both misfolded and properly folded, active (e.g., SCD1) ER proteins. Ube4A also controls expression of lipid uptake proteins such as CD36 and LDLR to reduce hepatocyte FFA levels. Together, protective functions of this E3 ubiquitin ligase reduce TAG formation, lipotoxicity and ER stress, delaying nutritional overload-induced development of hepatic steatosis, ER stress, steatohepatitis, and MASH (Figure 8 – top panel). In absence of hepatocyte Ube4A, the Ube4A-UBXN4 complex is disrupted, which impairs ERAD, enhancing protein misfolding-induced ER stress and SCD1-induced hepaticsteatosis. Moreover, FFA uptake is increased, which further disrupts hepatocyte protein and lipid homeostasis. Together, these pathogenic alterations enhance hepatic steatosis and ER stress and lipotoxicity-induced and liver injury, augmenting the development of steatohepatitis and MASH (Figure 8 – bottom panel).

**Figure 8.**
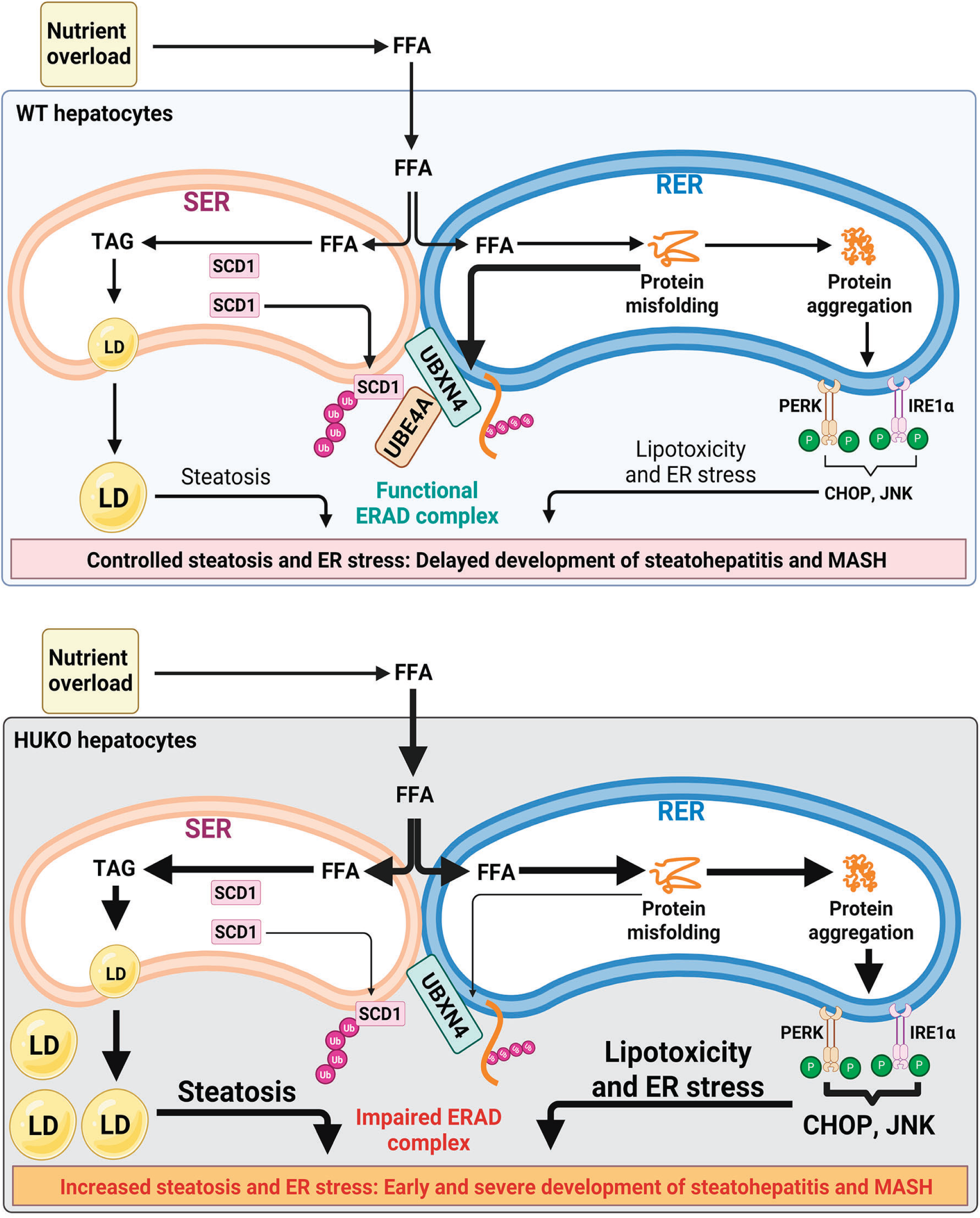
model: The hepatocyte Ube4A-UBXN4 complex enhances ERAD, which preserves ER lipid and protein homeostasis and protects against MASH development. Very thin, thin and thick arrows indicate reduced, normal and increased processes, respectively.

## Discussion

This study demonstrates that Ube4A mediates ERAD through its interaction with UBXN4, and downregulation and/or disruption of this protein complex augments MASLD and MASH development. This finding is novel as the roles of Ube4A and UBXN4 in MASLD and ER protein and lipid homeostasis had not been described. In fact, the impact of the cytosolic E3 ligases on hepatocyte-ERAD and MASLD are largely unknown except that whole body deletion of the U-Box E3 ligase CHIP causes age-induced hepatic steatosis and liver injury in chow-fed mice^56^.

The most robust effects of Ube4A deletion are on hepatocyte lipid accumulation and the associated pro-inflammatory and profibrotic stress response. This indicates that the progression of MASLD and development of steatohepatitis in these mice are consequences of hepatocyte lipid metabolic dysfunction in the setting of nutrient overload. In addition to increased TAG and DAG, altered levels of other lipids can also contribute to this phenotype. For example, PC and PE and the rate limiting enzymes in their biosynthetic pathway (choline-phosphate cytidylyl transferase A - Pcyt1A and ethanolamine kinase 2 - Etnk2) were significantly downregulated in HUKO livers. Glycerophospholipids, the major components of cell membranes, are also signaling mediators^57^. PC levels have been shown to be reduced in the liver samples of MASLD patients^57^. Reduced levels of PC activate Srebp1^58^, and trigger ER stress^59^, which can enhance MASLD and promote the development of steatohepatitis in HUKO mice. Whether Ube4A directly regulates glycerophospholipid levels by ubiquitinating PC/PE pathway enzymes will be determined in future studies.

Ube4A deletion increased or decreased many ER-processed proteins, suggesting that some of them are direct targets of the ligase and others could be altered secondarily. For example, Npc1, which undergoes substantial misfolding and ERAD^60^ is downregulated in HUKO livers, indicating Ube4A deletion further enhances misfolding and degradation of this protein by an Ube4A-independent ERAD machinery. This possibility is supported by downregulation of proteins involved in ER protein folding/processing and deubiquitination (e.g., Dnajc10, Creld2 and Usp19) and upregulation of other ERAD proteins (Ufd1, Erlin-2, Bag6 and Ube2b) in HUKO livers. Dramatic reduction of Npc1 in HUKO livers has disease relevance as genetic variation and heterozygosity of this gene are associated with obesity in humans and mice^61^. Whether Npc1 reduction is a direct consequence of Ube4A deletion or of increased MASLD in HUKO mice, needs further investigation.

Besides degrading misfolded proteins, ERAD also degrades properly folded, active lipid metabolic proteins such as SCD1^52^, ACSL3^62^, DGAT2^63^, APOB^64^, INSIG1^24^, HMGCR^65^, etc. SCD1 and ACSL3 are increased whereas INSIG1 and HMGCR are unaltered, indicating Ube4A deletion leads to the degradation of only a subset of the ERAD proteins. SCD1, a direct ERAD target of Ube4A, is the rate-liming enzyme of synthesis of MUFA, which are major substrates for the synthesis of TAG, phospholipids, and cholesteryl esters^51^. Accordingly, Scd1 deficiency favors lipid oxidation over synthetic pathways, leanness and improved insulin signaling^66,67^. Conversely, SCD1-mediated sequestration of FFAs into TAG/LDs may be a protective mechanism against the lipotoxicity of saturated fatty acids^68^. Nevertheless, inhibiting SCD1 has been shown to be beneficial in MASLD/MASH. The liver targeted SCD1 inhibitor aramchol (3β-Arachidyl amido cholanoic acid)^69^ reduces liver TAG^69,70^ and fibrosis in animal models of MASLD^71,72^ and alleviates various parameters of MASLD in human subjects (phase 2 clinical trials)^69,70,72–75^. Conceivably, SCD1 is one of the many ERAD targets of Ube4A to regulate the complex phenotypes of MASLD and MASH. Further studies are warranted to identify all the liver-specific EARD targets of Ube4A in healthy and MASLD livers.

Liver fibrosis, which was substantially increased in 32-week-GAN-D-fed female HUKO mice compared to LoxP controls was not measurably higher in 24-week-GAN-D-fed male HUKO mice despite showing significant upregulation in the fibrogenic pathway in both sexes of mice in absence of Ube4A. Although the exact reasons for these subtle sex-specific differences are unclear at this point, it should be noted that male HUKO mice developed MASLD features early (within 8-16-weeks of GAN-d feeding), which was maximal by 24 weeks. Possibly due to slower progression of these aberrations in female HUKO mice promoted increased levels of all the features of MASLD even after 32 weeks. Therefore, the timing of sex-specific MASLD development and the role of Ube4A in hepatic stellate cells (HSCs) could be determining factors, which will need to be further assessed in future studies.

Delayed insulin resistance in HUKO mice compared to UKO mice is explainable as Ube4A regulates insulin signaling in other metabolic tissues (e.g., skeletal muscle and adipose tissue)^37^. Interestingly, glucose uptake was substantially reduced in the BAT of both UKO (whole-body) and HUKO (hepatocyte-specific) mice, suggesting Ube4A regulates BAT and liver-BAT crosstalk to regulate insulin signaling and glucose homeostasis. A comparative metabolic study of adipocyte- and hepatocyte-specific Ube4A deletion will shed further light into this process in the future.

Ube4A-UBXN4-mediated ER homeostasis may play a role in protection from diseases. Downregulation of Ube4A and UBXN4 in human MASLD livers and disruption of the UBE4A-UBXN4 complex in the livers of MASLD mouse models strongly suggest that this novel mechanistic discovery has relevance in the development of MASLD. Ube4A also regulates obesity and glycemic profiles in mice^37^, and its downregulation is associated with obesity and type-2 diabetes^32–36^. Moreover, circulating autoantibodies against Ube4A correlate with disease severity in inflammatory bowel disease (IBD) patients^76,77^, and its expression is reduced in these patients^78^. Furthermore, UKO mice are sensitized to stressor-induced colitis^79^. UBXN4 is also strongly associated with metabolic diseases. It is downregulated in human MASL and MASH patients (GSE33814)^55^. The human *UBXN4* gene is located between two metabolic genes, miR-128-1 (thrifty gene)^80,81^ and lactase (*LCT*)^81^ in the locus (2q21.3). Moreover, FinnGen R11 dataset (>450,000 subjects) analysis identified UBXN4 among top 3 obesity-associated genes^82^. In conclusion, maintaining healthy levels of UBE4A and UBXN4 and their interaction may offer new therapeutic options for the treatment of obesity, MASLD and other diseases.

## Methods

### Human dataset analysis

#### RNA Microarray Data Analysis

Publicly available human liver RNA microarray datasets were analyzed to assess UBXN4 expression in control (N = 13) MASL (N = 19) and MASH (N = 12) subjects (included male and female subjects) using the GEO dataset GSE33814^55^. Expression matrices retrieved using GEOquery (v2.64.2). Expression values were log2-transformed when required. Differential expression analysis was performed using limma (v3.64.0). UBE4A and UBXN4 expression were visualized by plotting log2-transformed expression values for each dataset individually in Prism (v10.6.1).

### Generation and maintenance of HUKO mice

All animal studies were conducted in accordance with the protocols approved by the Institutional Animal Care and Use Committees (IACUC) of Saint Louis University and Vanderbilt University. All experimental groups consisted of age-matched littermates. Mice were maintained under controlled conditions (23^0^C, 12-hour light/dark cycles) with ad libitum access to standard chow diet and water and remained in good health throughout the duration of the studies.

The *Ube4A* gene is in the chromosome 9 of Mus musculus. Using CRISPR/Cas9 technology [33,34], two *LoxP* sites were inserted at the 5′- and 3′-ends of the last three exons, encoding the U-Box domain of Ube4A. The insertions were confirmed by next generation sequencing. Joining the 5′- and 3′-ends of the targeting locus generated two groups of C57BL6J mice: i) *Ube4A-Flox (LoxP* sites encompassing the targeting locus) and ii) *UKO* (complete deletion of the targeting locus of 9859 base pairs). Multiple lines of chimeric *UKO* mice were cross-bred with *WT* C57BL6J mice to obtain heterozygous *Ube4A-Flox* or *UKO* mice, which were inbred to generate respective homozygous lines.

Hepatocyte-specific *Ube4A* knockout (*HUKO)* mice were generated by crossing *Ube4A-floxed* (*LoxP*) mice with albumin-*Cre* transgenic mice (JAX: 003574) using a standard breeding protocol^83,84^. Genotypes were confirmed by PCR-based genotyping and validated via immunoblotting. Female *Ube4A-Flox* mice were bred with a male albumin-*Cre* transgenic mice. The resulting offspring (hets for both *Ube4A Flox* allele and *Alb-Cre* allele) were inbred to generate offsprings from which mice that were homozygous for *Ube4A-Flox* and heterozygous for *Alb-Cre* alleles were selected. These were further bred with *Ube4a-Flox/Flox* animals to generate cohorts with approximately 50% *Ube4a-Flox/Flox (LoxP)* & 50% *Ube4A-Flox/Flox + Alb-Cre/+* (HUKO) mice. Littermates of *LoxP* & *HUKO* animals were used for all experiments. Alb-Cre was genotyped using primers and protocol provided on the Jackson Laboratory website.

### Development of obesity in *WT* and *UKO* mice and obesity and MASLD in *LoxP* and *HUKO* mice

To induce obesity, 10-week-old, chow-fed male *WT* and *UKO* male mice were fed a HFD (TestDiet # 58Y1; fat 61.6 kcal%, carbohydrates 20.3 kcal %, and protein 18.1 kcal%) for the duration specified in the result section. To develop MASLD, 10-week-old, chow-fed *LoxP* and *HUKO* male and female mice were fed the Gubra-amylin-diet (GAN-D) (TestDiet # 5ZFL: fat 41.0 kcal% with 1.99% w/w cholesterol, carbohydrates 40.9 kcal% with 22.12% fructose, and protein 18.1 kcal%) for the durations specified in each experiment. After completion of the studies, mice were fasted for 5h followed by euthanasia by carbon dioxide asphyxiation.

### Body weight and composition

Body weight was measured between 10 - 11 AM by placing each mouse on a weighing scale. Body composition (fat, lean and fluid mass) was assessed using a Minispec LF-NMR (Brucker Optics) analyzer as previously described^37^.

### Hyperinsulinemic-euglycemic clamp

Sixteen-week-GAN-D-fed LoxP and HUKO and 12-week-HFD-fed WT and UKO (all male, littermates) were used for this study. Catheters were implanted into a carotid artery and a jugular vein of mice for sampling and infusions respectively five to seven days before the study^46^. Insulin clamps were performed on mice fasted for 5h using a modification of the original method^47^. After 3h fasting, an arterial blood sample was obtained to determine natural isotopic enrichment of plasma glucose. Ninety minutes prior to initiation of the clamp, a quantitative stable isotope delivery to increase glucose isotopic enrichment above natural isotopic labelling was initiated. [6,6-^2^H_2_]glucose was primed (16 mg) and continuously infused for a 90 min equilibration and basal sampling periods (0.8 mg/kg/min in saline). The insulin clamp was initiated at t=0min with a continuous insulin infusion (4 mU/kg body weight/min), which continued for 120 min. Arterial glucose was clamped using a variable infusion rate of glucose + [6,6-^2^H_2_]glucose (0.08 MPE), which was adjusted based on the measurement of blood glucose at 10 min intervals during the 120 min clamp period. By mixing the glucose tracer with the unlabeled glucose infused during a clamp, deviations in arterial glucose enrichment are minimized and steady state conditions are achieved. The calculation of glucose kinetics is therefore more robust^48^. Mice received heparinized saline-washed erythrocytes from donors at 5 μl/min to prevent a fall in hematocrit. Baseline blood or plasma variables were calculated as the mean of values obtained in blood samples collected at −15 and −5 min. At *t*=75 min, a continuous intravenous infusion of 2[^14^C]deoxyglucose ([^14^C]2DG) was initiated (0.32 µCi/min in saline). Blood was taken from 80–120 min for the determination of plasma [6,6-^2^H_2_]glucose enrichment and [^14^C]2DG. Clamp insulin was determined at *t*=120 min. At t=120 min, mice were anesthetized with pentobarbital (i.v.) and tissues were freeze-clamped for further analysis. Plasma insulin was determined by RIA. Plasma glucose enrichments ([6,6-^2^H_2_]glucose) were assessed by GC-MS as described previously^85^. Radioactivity of [^14^C]2DG in plasma samples, and [^14^C]2DG-6-phosphate in tissue samples were determined by liquid scintillation counting. Glucose appearance (Ra) and disappearance (Rd) rates were determined using steady-state equations^85^. Endogenous glucose appearance (endoRa) was determined by subtracting the GIR from total Ra. The glucose metabolic index (Rg) was calculated as previously described^86^.

### Glucose and insulin tolerance tests (GTT and ITT)

These studies were performed following our published protocols^37^. For GTT, mice were fasted overnight and received intraperitoneal (i.p.) injection of D-glucose (2 g/kg body weight). For ITT, 5h-fasted mice were injected intraperitonially with human insulin (0.5 U/kg body weight). Blood glucose levels were measured at baseline (0 min) and at designated time points (15, 30, 60, 90, 120 and 180 minutes post-injection for GTT and up to 120 minutes for ITT) via the tail-snip method using a calibrated glucometer.

### Whole-body energy expenditure

Mice were placed individually in metabolic cages at 23°C with a precise thermostatic control in a Comprehensive Laboratory Monitoring System (CLAMS; Columbus Instruments) and were acclimatized for 2 days. Afterwards, oxygen consumption, (VO_2_), carbon dioxide release (VCO_2_), respiratory exchange ratio (RER), Energy expenditure (EE) and locomotor activity were measured for 7 days^37,83^.

### Tunicamycin-Induced ER Stress in Mice

LoxP and HUKO mice were given a single intraperitoneal injection of tunicamycin (1 mg/kg body weight, delivered in 200 µL of 150 mM dextrose). The control animals received an equivalent volume of vehicle solution (150 mM dextrose). Following treatment, animals were monitored daily, and tissues were harvested on day 5 for subsequent analyses.

### Tissue collection

At the conclusion of the studies, mice were fasted for 5h (to avoid munching-related spikes in metabolic parameters) and euthanized. Blood was collected, serum separated by centrifugation and stored at -80^0^C for downstream analyses. The liver tissues were weighed, flash-frozen and stored at -80^0^C for immunoblotting, qRT-PCR, lipidomics, and proteomics studies. A portion of the liver was fixed in 10% neutral-buffered formalin at 4°C for histological evaluation.

### Measurement of TAG from serum and liver

Serum levels of TAG were analyzed using commercial assay kits following manufacturers’ instructions. For serum analyses, samples were mixed with the provided reagents in a microplate reader for appropriate time, and absorbance of the colorimetric product was measured at 540nm. Concentrations (mg/dL) were determined from the standard curve. Liver TAG was measured in total lipids, extracted from the tissue, following the above-mentioned protocols. Levels were normalized to liver weight (mg/g).

### Serum levels of ALT and AST

ALT and AST were measured using commercial kits in accordance with the manufacturer’s instructions. After incubating serum samples with the respective substrate reagents, the enzymatic reactions were carried out at 37°C and absorbance was monitored at 340 nm using a microplate reader. Enzyme activities were calculated according to the kit instructions and reported in units per liter (U/L).

### Hematoxylin and Eosin (H&E) staining and MASLD feature scoring

Fixed liver tissues were processed, paraffin-embedded, and sectioned at 4 µm thickness. Sections were deparaffinized, rehydrated, and stained with H&E following standard histological procedures at the core facility. Images were captured at 10X magnification with 100 µm scale bar to show the lobular distribution of macrovesicular steatosis. Inflammation was assessed at higher magnification (20X), with a 100 µm scale bar applied to facilitate identification and localization of inflammatory loci.

MASLD features were assessed using a validated semiquantitative scoring system for rodent models^40^. Hepatic macrosteatosis, microsteatosis and hepatocellular hypertrophy were graded on a scale from 0 to 3, corresponding to the percentage of affected hepatocytes: 0 (<5%), 1 (5–33%), 2 (34–66%), and 3 (>66%). Lobular inflammation was evaluated by quantifying the number of inflammatory foci per 200X microscopic field. The data presented represents the mean scores from 8 mice per experimental cohort.

### PicroSirius red staining and analysis of quantification of total collagen content

Collagen deposition was assessed using the Picrosirius Red Stain kit following the manufacturer’s protocol. Briefly, liver tissue sections were incubated in Picrosirius Red solution. Following staining, slides were then washed with distilled water, dehydrated with ethanol, cleared and mounted. Sections were imaged at 100× magnification, with a 100 µm scale bar under bright field microscope. Collagen deposition was assessed by morphometric analysis as previously described^37^. Briefly, the collagen-positive areas were calculated as a percentage of the total tissue area using digital image analysis with the ImageJ software (NIH). For comparative analysis, the collagen contents in HUKO mouse livers were normalized to the corresponding LoxP control group.

### Immunohistochemical staining for CD11C and α-SMA

Immunohistochemical staining for CD11C and α-SMA was performed on an automated staining platform using a DAB chromogenic detection system, in accordance with standard procedures at the institutional core facility. Briefly, fixed liver sections were incubated with primary antibodies for overnight at 4°C to detect CD11C (1:50 dilution,) and α-SMA (1:150 dilution). After washing with PBS, slides were incubated with respective anti secondary antibody. DAB (diaminobenzidine) was used as a chromogen for visualization. The slides were then counterstained with hematoxylin, dehydrated, cleared, and cover slipped. Finally, slides were examined under microscope, and images were acquired at 10X magnification with 50 um scale bars. Brown staining was considered indicative of positive expression.

### RNA Extraction and qRT-PCR

To measure gene expression levels in the liver samples, qRT-PCR studies were performed following a standardized protocol as previously described^37^. Briefly, total RNA was isolated from frozen liver tissues using Trizol reagent according to manufactures instructions. The extracted RNA was subsequently purified using RNeasy mini kit columns from Invitrogen. Following purification, RNA concentration was assessed from spectrophotometrically and 2ug of total RNA was reverse-transcribed into cDNA using a high-capacity cDNA reverse transcription kit. Quantitative real-time PCR (qRT-PCR) was performed using SYBR Green-based detection on a QuantStudio real-time PCR system. The relative fold change in gene expression levels was analyzed using the comparative 2^−ΔΔCT^ method, normalized to housekeeping genes such as *RPLP0*. For primary hepatocytes, media was aspirated and cells collected in 1mL TRI reagent (T9424-200ML, Sigma-Aldrich, Missouri, USA) for total RNA isolation per manufacturer’s instructions. After RNA quantification, 500 ng total RNA was reverse transcribed to cDNA by PR1MA qMAX cDNA synthesis kit. qRT-PCR was performed using PR1MA qMax Green low rox master mix on a QuantStudio real-time PCR system. The relative levels of gene expression were normalized to 36b4/*Rplp0* as housekeeping control using the 2^−ΔΔCT^ method.

### Proteomic studies for relative abundance of liver proteins in HUKO and LoxP mice

Equal (20 μg) amount of total protein was for digested (with trypsin/lys C). The MS analysis of digested proteins was immediately performed to ensure high-quality tryptic peptides with minimal non-specific cleavage. Nano-liquid chromatography tandem mass spectrometry (Nano-LC/MS/MS) was performed on a Thermo Scientific Q Exactive HF Orbitrap mass spectrometer equipped with an EASY Spray nanospray source (Thermo Scientific) operated in positive ion mode. The LC system was an UltiMate™ 3000 RSLCnano system from Thermo Scientific. The MS/MS was acquired according to standard conditions. All MS/MS spectra were analyzed using the Chimerys node (Thermo Fisher Scientific, San Jose, CA, USA; Proteome Discoverer 3.2.0.450). Precursor ion intensity label free quantitation was done using Proteome Discoverer (Thermo Fisher Scientific vs 3.2.0.450).

The two groups were compared using a “non-nested” study factor. Normalization was derived by using all peptides. Protein abundances were calculated by summed abundances, meaning the protein abundances are calculated by summing sample abundances of the connected peptide groups. Fisher’s exact test (pairwise ratio-based) was used to calculate p-values with low intensity resampling value imputation included. Adjusted p-values were calculated using Benjamini-Hochberg.

Proteins were considered differentially expressed if adjusted p-value ≤ 0.05 and an absolute fold change ≥ 1.5. KEGG and Gene Ontology (GO) pathways were determined to be enriched using limma (v3.64.0) on Entrez Gene IDs corresponding to genes encoding the differentially expressed proteins. Dot plots of top enriched pathways were generated in Python using matplotlib (v3.10.1) and seaborn (v0.13.2). Heatmaps were created using normalized abundance scores for each sample. A pseudocount of half the minimum non-zero value present in heatmap was added to all values. Differentially expressed proteins with an adjusted p-value ≤ 0.05 were plotted after calculating a z-score per protein across samples using Prism (v10.6.1). Volcano plots were generated in R using ggplot2 (v4.0.2) by plotting log2 fold change on the x-axis against −log10(p-value) on the y-axis.

### Proteomic studies to identify the Ube4A interacting proteins in the liver

The Ube4A-interactome from liver was identified using our published methods^37,84^. Liver samples from Human MASH- and mouse-livers (chow- and 18-week-Western diet-fed (samples were taken from our publication^84^) were used for these studies. Equal (2 mg) amounts of protein were used for immunoprecipitation (after preclearing with Ig-control) with Ig-control or Ig-Ube4A antibodies. Proteins co-precipitated with both antibodies were identified by LC-MS/MS and proteins that are specifically coprecipitated with Ube4A were considered as putative Ube4A interactors (numbers denote protein spectra; none detected in Ig-control).

### Lipidomics for relative abundance of liver lipids in HUKO and LoxP mice

Liver tissue portions (100-200 mg) were homogenized in 300 µL 50:50 (v:v) water:methanol using zirconium beads on a BeadBug homogenizer (Benchmark). Homogenate (10 mg) was combined with 1.4 mL 2:1:1.5 chloroform:methanol with 1 mM dibutylhydroxytoluene (BHT): water and 5 µL SPLASH LIPIDOMIX internal standard (Avanti), sonicated 15 minutes, and incubated at -20 °C overnight. The organic layer was removed via centrifugation and a second extraction was performed using an additional 1.4 mL 2:1:1.5 chloroform:methanol with 1 mM BHT:water. The extraction was vigorously mixed and incubated at -20 °C for 1 hour. The organic layers were combined and dried under vacuum to completeness. Samples were reconstituted in 500 µL 90:10 methanol:chloroform. A portion (50 µL) was used for total lipid pre-quantitation while the remainder was analyzed by LC-MS/MS. Total lipid pre-quantitation was performed using the sulfo-phospho-vanillin assay (SPVA) against a total lipid liver calibrant (0-2000 µg/mL; Avanti Lipids)^1^.

Liquid chromatography (LC) was performed using a Thermo Fischer Scientific UltiMate 3000 RSLCnano system equipped with degasser, pump, column compartment, and autosampler operating at a column temperature of 40 °C and autosampler temperature of 10 °C. Sample loading was normalized to 3 µg total lipid and separated on an Acquity UPLC CSH C18 column (1.0 x 100 mm, 1.7 µm, 130 Å, Waters) operating at 80 µL/min flow rate and 65 °C column temperature using a 10 minute gradient from 50% to 99%B. Separation was performed using mobile phases (A) 60/40 (v/v) acetonitrile/water and (B) 90/8/2 (v/v) isopropanol/acetonitrile/water, both contained 10 mM ammonium formate and 0.1% formic acid. Mass spectrometry (MS) was performed on a Burker Daltonics TIMStof Pro 2 equipped with a heated electrospray ionization source operating in positive mode using a 100 ms TIMS ramp rate. Parallel accumulation and serial fragmentation (PASEF) were applied to ions 0.55-1.90 V*s/cm2, m/z 100–1350 mass range using an m/z 2 mass window and 30 eV collision energy. Active exclusion was engaged after a single spectrum for 0.1 minutes unless the current-to-previous intensity ratio was greater than or equal to 3.0. A pooled QC sample was used to monitor instrument performance throughout the sequence. Mass and mobility accuracy was maintained using a 90:10 (v:v) ESI tune mix (Agilent):10 mM sodium formate solution mixture. Processing was performed in Metaboscape (v.2026 Bruker Daltonics) using Lipid Blast (v66)^2^ and MSDial^3^ spectral libraries followed by manual spectral evaluation.

### Immunoblotting studies

Immunoblotting was performed according to our previously published protocols^37^. Briefly, frozen liver tissues were homogenized in lysis buffer containing 20 mM Tris-HCl (pH 7.4), 150 mM NaCl, and 1% Triton X-100, supplemented with protease and phosphatase inhibitor cocktails. Homogenates were incubated on ice for 30 minutes and then centrifuged at 12,000 × g for 15 minutes at 4°C. The supernatants were collected, and total protein concentration was determined using a BCA protein assay kit. Equal amounts of protein (20 μg per sample) were separated on 10% SDS-PAGE gels and transferred to PVDF membranes. Membranes were blocked in 5% non-fat dry milk in TBST for 1 hour at room temperature and incubated overnight at 4°C with primary antibodies against target proteins. After washing, membranes were incubated with appropriate HRP-conjugated secondary antibodies and developed using enhanced chemiluminescence (ECL) reagents. Protein bands were visualized using a Bio-Rad Chemidoc imaging system and quantified by using ImageJ, respectively. Densitometric quantification of band intensity was performed using ImageJ software (NIH), and target protein expression was normalized to loading controls such as GAPDH.

### Primary hepatocyte isolation and hepatocyte TAG measurement

Mice were anesthetized with 3% isoflurane and then euthanized via cervical dislocation. Quickly the abdominal cavity was exposed, and primary murine hepatocytes were isolated by perfusion of the portal vein in situ with HBSS then 1 mg/mL collagenase digestion as previously performed^37,87^. Digested liver solution was passed through a 100 μm cell strainer, then isolated hepatocytes were washed with 10%FBS-DMEM twice, counted by hemocytometer, and plated onto collagen-coated 6-cm dishes or 6-well culture plates at 250,000 live cells/mL. Three hours after plating, cells were washed with PBS and placed into serum-free high-glucose DMEM (Sigma) with vehicle (DMSO) or the SCD1 inhibitor (CAY10566: 4 μM) for 20 hours. Afterwards, media was removed and cells scraped in PBS. Cells were then pelleted by centrifugation and suspended in PBS (150 μL). Cell suspension was diluted 1:1 with 1% sodium deoxycholate to extract lipids. TAG was measured by commercial colorimetric assay. Protein content was measured by BCA to normalize TAG.

### Generation of mutants of human UBE4A

For the generation of mutants (described in results section), we designed mutagenic primers introducing the desired deletions in the target sequence. We performed the PCR amplification using wild-type genes as a template and the mutagenic primers with SalI and NotI restriction sites. The cDNA was subcloned into a bacterial expression vector PCMV (GST-tagged). After sequence confirmation, GST tagged PCMV plasmids were transformed into the E. Coli strain BL21 (DE3). Transformed colonies were selected on antibiotic containing LB agar plates and grown overnight. Plasmid DNA was extracted from individual colonies, and the presence of desired deletion was confirmed by sanger sequencing.

### Cell culture experiments

Huh-7 and HEK293 cells were cultured in Dulbecco’s modified Eagle’s medium (DMEM) supplemented with 10%FBS, 100 U/mL penicillin, and 100 μ g/mL streptomycin at 37°C with 5% CO_2_. HEK 293 cells were transiently transfected with 2ug of expression plasmids using the Polyplus Jetprime transfection reagent according to the manufacturer’s instructions. Post 5h transfection, culture media containing transfection reagent was replaced with fresh medium. After 48h, cells were lysed for downstream analyses.

Huh-7 cells were transiently transfected with the control- and UBE4A-overexpressing plasmids using lipofectamine-3000. Media was replaced with fresh complete growth media 16h post-transfection. After 48h, cells were lysed for downstream analyses.

For shRNA delivery, 60-70% confluent Huh-7 cells were transduced with adenoviral vectors expressing control- and UBE4A-shRNA (MOI-1.75×10^8^ PFU/ml). After 16h, media was replaced with fresh complete medium. After 72h, cells were lysed for downstream analyses.

### GST-pull down assays in HEK293 cells

HEK293 cells expressing GST-UBE4A and its deletion mutants were harvested 48h after expression. Cells were lysed, protein concentrations were measured, and lysates containing equal concentration (1 mg) of total proteins were incubated with glutathione-agarose beads for 16 hours (overnight) at 4 °C with gentle rotation to allow binding of the GST-tagged protein to the beads. After incubation, the beads were washed three times with washing buffer for 15 min each with agitation to remove unbound proteins. The bead-bound GST-tagged protein and its interacting proteins were eluted from beads and analyzed by SDS-PAGE, followed by immunoblotting studies.

### Immunoprecipitation studies

For immunoprecipitation studies, approximately 2mg of protein lysate were precleared with IgG Control and Protein A/G agarose beads. Precleared lysates were then incubated with protein A/G-agarose beads along with either IgG control or anti-UBE4A antibodies for 16 hours (overnight). Afterwards, the beads were collected by centrifugation at low speed at 4°C and washed (3X, 15 minutes each) with a wash buffer (lysis buffer without the protease and phosphatase inhibitors). Bound proteins were eluted by boiling the beads in 2X loading buffer.

### Immunofluorescence

HEK293 Cells were transfected with plasmid constructs of human UBE4A and UBXN4 and UBE4A mutants (as described in results section) for 48 hours, followed by immunostaining protocol. For immunostaining cells were fixed in 4% PFA for 15 minutes, permeabilized in ice-cold 100% Methanol at -20°C for 10 minutes, blocked for 1 hour, incubated in primary antibody solution at 4°C overnight & secondary antibody solution at room temp for 1 hour and DAPI solution in PBS for 15 with 3X PBS washes between every step mentioned. Primary antibody dilutions: Anti-Myc Tag Mouse mAb (1:1000), Anti-HA Tag Rabbit mAb (1:1000), Anti-HA Tag Mouse mAb (1:100), Anti-Calnexin Rabbit antibody (1:100). Secondary antibodies: Alexa Fluor 647 Goat Anti-Mouse (1:1000), Alexa Fluor 647 Goat Anti-Rabbit (1:1000), Alexa Fluor 488 Goat Anti-Mouse (1:1000). Slides were imaged using a Leica TCS SP5 Confocal microscope with a 63X water objective and laser lines UV 405nm, Argon 488nm & HeNe 633nm.

### Quantification and statistical analysis

All data are presented as ± Standard Error of the Mean (SEM). Animals showing any signs of sickness were excluded from the study. The number of mice (n) used in each experiment is indicated in the figure legends. Statistical difference between two groups was determined by unpaired 2-tailed Student’s *t* test (with Welch’s correction). For multiple groups comparison, 2-way ANOVA followed by Sidak’s multiple comparison test was used. ImageJ software (NIH) was used to quantify the western blots. All statistical analysis was performed using Graphpad prism (Version 10). The result was considered as statistically significant if the P value is less than 0.05. Schematic images were created using Bio render.

## Acknowledgement

This research was funded by the NIH grants R01 DK132162 and startup funds from University of Florida to Anutosh Chakraborty; NIH R01 DK132162, R35 GM159817 American Heart Association 24TPA1299435 to Kyle S. McCommis. Hyperinsulinemic-euglycemic clamp studies were performed by the Vanderbilt Mouse Metabolic Phenotyping Center (DK020593, DK135073). The Vanderbilt Hormone Assay and Analytical Core performed the insulin analysis during clamp (DK020593). Relative abundance proteomics and lipidomics studies were performed at the UF Mass Spectrometry Research and Education Center by Drs. Kari Basso and Laura Bailey, and Katie Heiden (NIH S10 OD021758-01A1 and NIH S10 OD030250-01A1). The UBE4A-interactome studies were performed at the Washington University Proteomics Shared Resource (WU-PSR) (R. Reid Townsend MD. Ph.D., director). The expert technical assistance of Petra Erdmann-Gilmore, Yiling Mi, and Rose Connors is gratefully acknowledged. WU-PSR is supported in part by the WU Institute of Clinical and Translational Sciences (NCATS UL1 TR000448), the Mass Spectrometry Research Resource (NIGMS P41 GM103422), and the Siteman Comprehensive Cancer Center Support Grant (NCI P30 CA091842). Histology and IHC studies were performed at the Molecular Pathology laboratory at UF and the histology core facility at SLU. Technical assistance from Kjirstin Osland, Lourdes Hindi, Shreya Das and Diya Patel are acknowledged. We also thank Sandip Mukherjee, Naomi Msengi and Jake Haubner for sharing reagents etc. We thank Kathleen Donovan and colleagues from Saint Louis University Comparative Medicine for helping with mouse colony maintenance. We also thank the Burris lab for helping with reagents and equipment. The schematic models were created with Biorender.com

## Author contribution

A.C. conceived, outlined, and directed the project and provided critical input throughout the study; M.C. generated *HUKO* mice and maintained mouse colonies; N.L.C., V.B. and M.C. performed majority of the experiments; L.L. performed hyperinsulinemic-euglycemic clamps, M.H. performed pathway analysis for proteomics studies and secondary analysis of published human MASLD/MASH datasets; I.R. assisted in analysis of human MASLD/MASH datasets; G.K. and K.S.M. shared reagents and performed experiments in primary hepatocytes; C.W.V.K. performed Alphafold3 studies; D.C. scored histology slides for MASLD parameters; S.K. quantified PicroSirius stained-slides for fibrosis; A.P., S.P., K.M., and S.G. assisted with construct generation, biochemical experiments and genotyping; B.A.T. provided inputs throughout the study; A.C., N.L.C. analyzed data and generated most figures; M.H., G.K., K.S.M. and C.W.V.K. also analyzed data and generated figures for their part of the study; A.C. and N.L.C. wrote the first draft; all the authors read, edited and approved it; A.C. submitted it to the journal.

## Declaration of competing interest

B.T. has been an advisor or consultant for Alimentiv, Allergan, Allysta, Alnylam, Amgen, Arrowhead, Axcella, Boehringer Ingelheim, BMS, Coherus, Cymabay, Enanta, Fortress, Genfit, Gilead, High Tide, HistoIndex, Innovo, Intercept, Ionis, LG Chem, Lipocine, Madrigal, Medimmune, Merck, Mirum, NGM, NovoNordisk, Novus Therapeutics, pHPharma, Sagimet, Target RWE, 89Bio; he has stock options in HepGene; he has institutional research grants from Allergan, BMS, Cirius, Enanta, Genfit, Gilead, Intercept, Madrigal, NGM. A.C. is a co-founder of InoKare Therapeutics. The other authors do not have any conflict of interest.

## Supplementary Figure legends

**Figure S1: Hepatocyte-specific Ube4A deletion augments GAN diet-induced steatosis, inflammation and injury of the liver in male mice.** A. Serum TAG levels in 8- and 16-week-GAN-D-fed male LoxP and HUKO mice.

B-F. Oxygen consumption (VO₂), energy expenditure (EE), respiratory exchange ratio (RER), ambulatory activity (Xamb), and total activity (Xtot) in LoxP and HUKO mice following 6 weeks of GAN-D diet feeding.

G-H. Relative expression of fibrotic marker genes in the livers of 16- and 24-week-GAN-D-fed male LoxP and HUKO mice (N=6/cohort).

I-J. Representative Picrosirius Red–stained liver sections along with the quantification from 24-week-GAN-D-fed male LoxP and HUKO mice. Magnification - 10X, scale bar - 100 µm.

K-L. Body weight, fat mass and liver weight in 24-week-GAN-D-fed male mice (N=8/cohort).

M. Liver TAG levels in GAN-D diet fed male mice (N=8/cohort).

N. Serum TAG levels in 24w GAN-D diet–fed male mice (N=8/cohort).

**Figure S2: GAN-D-induced MASLD features are amplified in female HUKO mice** A-C. Body weight, and fat and lean mass in 32-week-GAN-D-fed female LoxP and HUKO mice (N = 8 per cohort).

**Figure S3: GAN-D-fed male HUKO mice exhibit late-onset insulin resistance** A-B. Extension of Figure 3A. Fasting glucose and plasma insulin levels during clamps in 16-week-GAN-D-fed male LoxP and HUKO mice.

C. Extension of Figure 3D. Tissue-specific 2-DG uptake (Rg) in metabolic tissues of LoxP and HUKO mice.

D-E. Raw values of GTT and ITT in 12-13-week-GAN-D-fed male LoxP and HUKO mice (n = 8/cohort).

F-G. Fasting glucose and plasma insulin levels during clamps in 12-week-HFD-fed male WT and UKO mice (N=6 – WT and 9 – UKO).

H-I. GIR and endoRa values in 12-week-HFD-fed male WT and UKO mice.

J-K. Rd values and insulin-induced shifting in Rd values in WT and UKO mice. L-M. Tissue-specific 2-DG uptake (Rg) in various tissues of WT and UKO mice.

**Figure S4: Hepatocyte-Ube4A deletion upregulates lipid accumulation pathways, leading to increased TAG accumulation and hepatic steatosis in GAN-D-fed mice** A-B. Extension of Figure 4A. Lipid class distribution in 16-week-GAN-D-fed male LoxP and HUKO liver samples by mass spectrometry (N = 5 per cohort).

C. PCA plots of lipidomic study.

D. Volcano plots showing relative levels of various species of TAG and DAG in the livers of male LoxP and HUKO mice.

E. Levels of various species of TAG and DAG in male LoxP and HUKO livers.

F. Extension of Figure 4G. Densitometric quantification of lipogenic proteins in LoxP and HUKO mice.

G. Relative hepatic levels of lipoprotein gene expression in 16-week GAN-D-fed male mice (N = 6 per cohort).

**Figure S5: The liver proteome of GAN-D-fed HUKO mice exhibits alterations in MASLD, ER processing and ERAD pathway proteins.** A. Extension of Figure 5A. Volcano plot, summarizing the altered proteins in HUKO livers.

B-F. Tables representing examples of proteins from certain pathways that are altered in HUKO livers.

**Figure S6: Hepatocyte ER stress is increased in absence of Ube4A** A. Relative hepatic expression of ER stress genes Bip, Xbp1, Ire1a and eIF2s1a in 24-week-GAN-D-fed male LoxP and HUKO mice (N=6/cohort).

**Figure S7: Interaction with the ERAD mediator UBXN4 facilitates ER-localization of Ube4A and degradation of Scd1 in the liver** A-B. Extension of Figure 7A. Peptides of Ube4A and UBXN4 (highlighted), identified from chow-fed WT mouse livers by LC-MS/MS.

C. Immunoprecipitation of UBXN4 co-immunoprecipitated Ube4A in chow fed mice liver.

D. Identification of endogenous UBXN4 and Ube4A in human embryonic kidney (HEK293) cells by Ube4A-immunoprecipitation followed by LC-MS/MS.

E. Alphafold3 prediction of a very strong and specific interaction between the N-termini of human Ube4A and human UBXN4.

F. GST-pulldown assay in HEK293 cells using wt-, N-terminal 1-47 amino acids deleted (d-1-47)-, F15A-, F15S- and DM (F15A-L18A)-UBE4A for endogenous UBXN4 interaction.

