## Supplemental Figures for "Hepatocyte UBE4A protects against MASLD and MASH by maintaining ER homeostasis via UBXN4 interaction"

**Figure S1: Hepatocyte-specific *Ube4A* deletion augments GAN diet-induced steatosis, leading inflammation and injury of the liver in male mice**

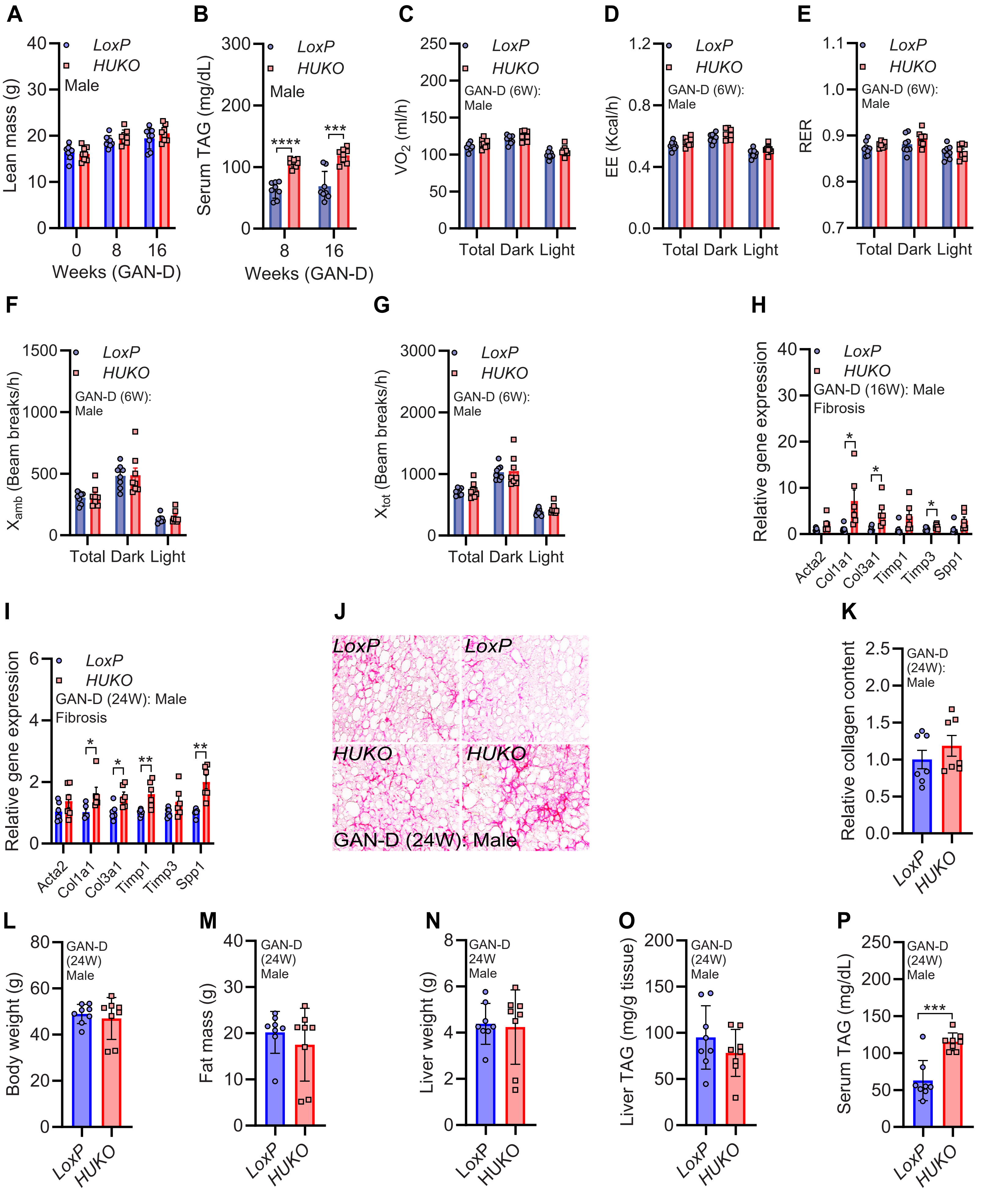

**Figure S2: GAN-D-induced MASLD features are enhanced in female HUKO mice**

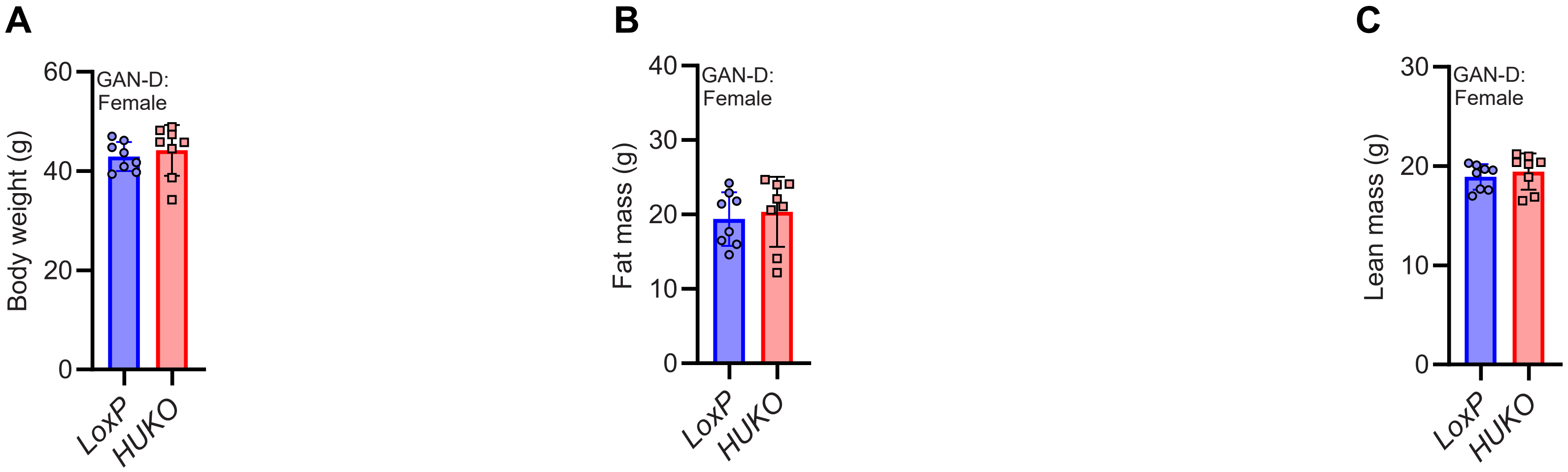

### Figure S3: GAN-D-fed HUKO mice exhibit late-onset insulin resistance

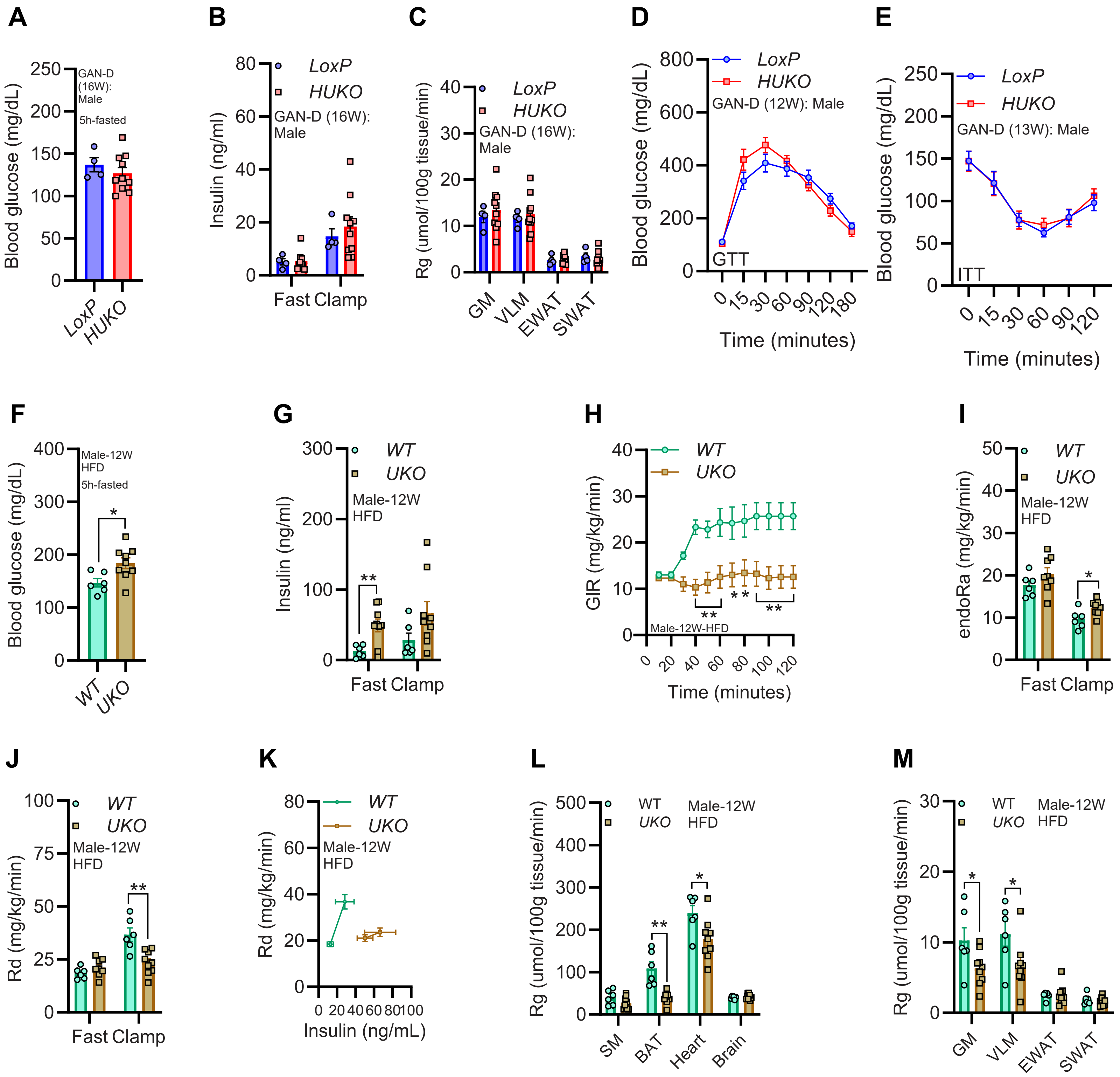

**Figure S4: Hepatocyte-Ube4A deletion upregulates lipid accumulation pathways, leading to increased TAG accumulation and hepatic steatosis in GAN-D-fed mice**

**A**

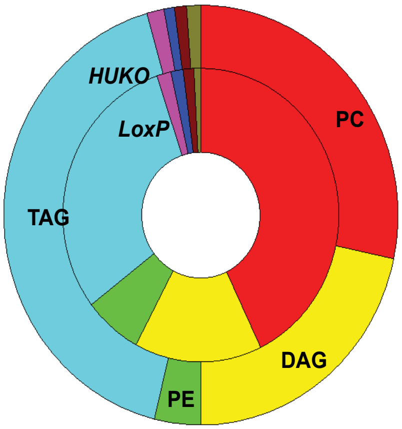


**B**

| Lipid class | *HUKO* | *LoxP* |
|---|---|---|
| Phosphocholine | 29% | 43% |
| Diacylglycerol | 22% | 15% |
| Phosphoethanolamine | 4% | 7% |
| Triacylglycerol | 42% | 31% |
| Ceramide | 1% | 2% |
| Sphingomyelin | 0.9% | 1.5% |
| LysoPhosphocholine | 1.0% | 1.3% |
| Phosphoserine | 0.14% | 0.06% |
| Cardiolipin | 0.10% | 0.08% |
| Cholesterol ester | 0.5% | 0.3% |
| LysoPhosphoethanolamine | 0.1% | 0.2% |
| Phosphoinositol | 0.07% | 0.08% |
| Quinone | 0.2% | 0.1% |
| Fatty amide | 0.02% | 0.01% |
| Glycerophosphate | 0.007% | 0.013% |
| Isoprenoid | 0.003% | 0.004% |
| Secosteroid | 0.006% | 0.007% |

**C**

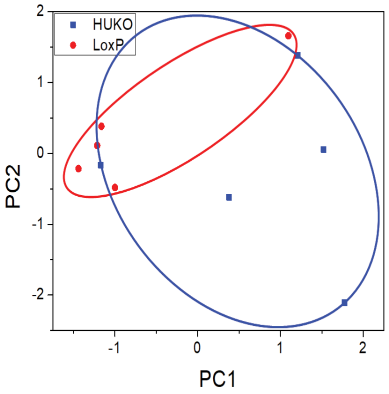


**D**

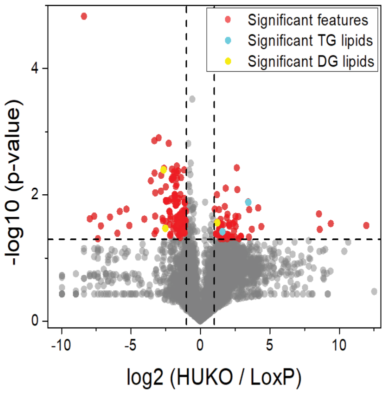


**E**

| Lipid | HUKO/LoxP (fold) | P values | Significance |
|---|---|---|---|
| TAG, tripalmitin (16:0_16:0_16:0) | 10.944 | 0.01699 | * |
| TAG (14:1_16:1_18:1) | 3.002 | 0.038153 | * |
| DAG (16:0_18:1) | 2.312 | 0.027451 | * |
| DAG (15:0_16:1) | 4.782 | 0.054321 | No |
| TAG (14:0_16:1_16:1) | 4.098 | 0.056821 | No |
| TAG (14:0_14:1_16:0) | 5.502 | 0.070919 | No |
| TAG (16:1_18.2_20:4) | 7.544 | 0.071477 | No |

**F**

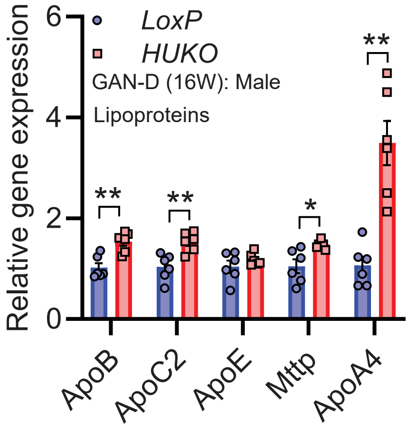

#### Figure S5: The liver proteome of GAN-D-fed HUKO mice exhibits alterations in MASLD, ER processing and ERAD pathway proteins

**A**

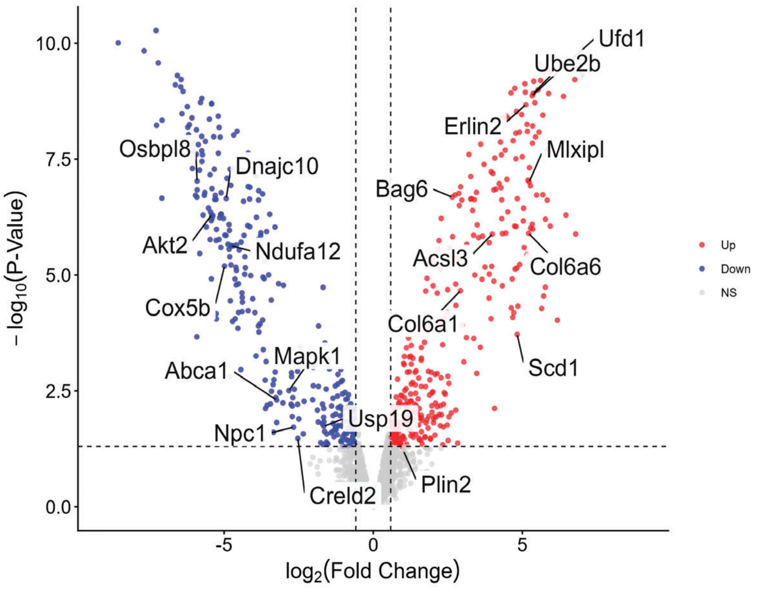


**B**

| Lipogenesis and lipid/cholesterol metabolism | | |
|---|---|---|
| **Examples of upregulated proteins** | **Symbol** | **Function** |
| Carbohydrate-responsive element-binding protein | Mlxipl/ChREBP | Transcription |
| Fatty acid CoA ligase | Acsl3 | Lipogenesis |
| Acyl-CoA desaturase 1 | Scd1 | Lipogenesis |
| Perilipin-2 | Plin2 | LD stability |
| **Examples of downregulated proteins** | | |
| NPC intracellular cholesterol transporter 1 | Npc1 | Lipid export |
| Phospholipid-transporting ATPase ABCA1 | Abca1 | Lipid export |
| Cytochrome c oxidase subunit 5B, mitochondrial | Cox5b | OxPhos |
| NADH dehydrogenase [ubiquinone] 1 alpha subcomplex subunit 12 | Ndufa12 | OxPhos |
| 2-methoxy-6-polyprenyl-1,4-benzoquinol methylase, mitochondrial | Coq5 | OxPhos |

**C**

| Insulin signaling | | |
|---|---|---|
| **Examples of downregulated proteins** | **Symbol** | **Function** |
| Mitogen-activated protein kinase 1 | Mapk1 | Enhances insulin signaling |
| DCC-interacting protein 13-alpha | Appl1 | Enhances insulin signaling |
| Syntaxin-4 | Stx4 | Enhances insulin signaling |
| Starch-binding domain-containing protein 1 | Stbd1 | Enhances insulin signaling |
| RAC-beta serine/threonine-protein kinase | Akt2 | Enhances insulin signaling |
| Flotillin-1 | Flot1 | Enhances insulin signaling |

**D**

| Fibrosis | | |
|---|---|---|
| **Examples of upregulated proteins** | **Symbol** | **Function** |
| Collagen alpha-6(VI) chain | Col6a6 | Marker of fibrosis |
| Collagen alpha-1(VI) chain | Col6a1 | Marker of fibrosis |
| Fibrillin-1 | Fbn1 | Marker of fibrosis |
| TGF beta-induced protein ig-h3 | Tgfbi | Activates HSC |
| Periostin | Postn | Activates HSC |
| Decorin | Dcn | Inhibits TGF beta |
| **Examples of downregulated proteins** | | |
| EMILIN-1 | Emilin1 | Antifibrotic |
| Integrin beta-3 | Itgb3 | Fibrotic |

**E**

| Proteins that mediate ERAD | | |
|---|---|---|
| **Examples of upregulated proteins** | **Symbol** | **Function** |
| Ubiquitin recognition factor in ERAD 1 | Ufd1 | ERAD receptor |
| Erlin-2 | Erlin2 | ERAD receptor |
| Large proline-rich protein BAG6 | Bag6 | Cytosolic ERAD chaperone |
| Golgi to ER traffic protein 4 homolog | Get4 | Bag6 cofactor |
| Ubiquitin-conjugating enzyme E2 B | Ube2b | E2 conjugation enzyme |
| **Examples of downregulated proteins** | | |
| DnaJ homolog subfamily C member 10 | Dnajc10 | ER chaperone |
| Protein disulfide isomerase | Creld2 | ER chaperone |
| Ubiquitin carboxyl-terminal hydrolase 19 | Usp19 | ER resident deubiquitinating enzyme |

**F**

| Proteins that undergo ERAD | | |
|---|---|---|
| **Examples of upregulated proteins** | **Symbol** | **Function** |
| Fatty acid CoA ligase | Acsl3 | Lipogenesis |
| Acyl-CoA desaturase 1 | Scd1 | Lipogenesis |
| Syntaxin-18 | Stx18 | Golgi to ER traffic |
| Gap junction beta-1 protein | Gjb1 | Gap junction |
| Fibrillin-1 | Fbn1 | ECM |
| **Examples of downregulated proteins** | | |
| NPC intracellular cholesterol transporter 1 | Npc1 | Lipid export |
| Phospholipid-transporting ATPase ABCA1 | Abca1 | Lipid export |
| Oxysterol-binding protein-related protein 8 | Osbpl8 | Lipid transport |
| Ras-related protein Rab-8A | Rab8a | Membrane trafficking |
| Nicastrin | Ncstn | Membrane protein regulation |

**Figure S6: Hepatocyte ER stress is increased in absence of Ube4A**

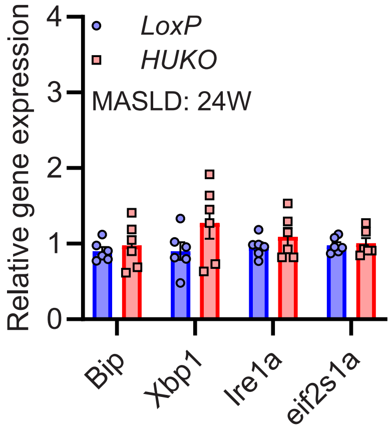

### Figure S7: Interaction with the ERAD mediator UBXN4 facilitates ER-localization of Ube4A and degradation of Scd1 in the liver

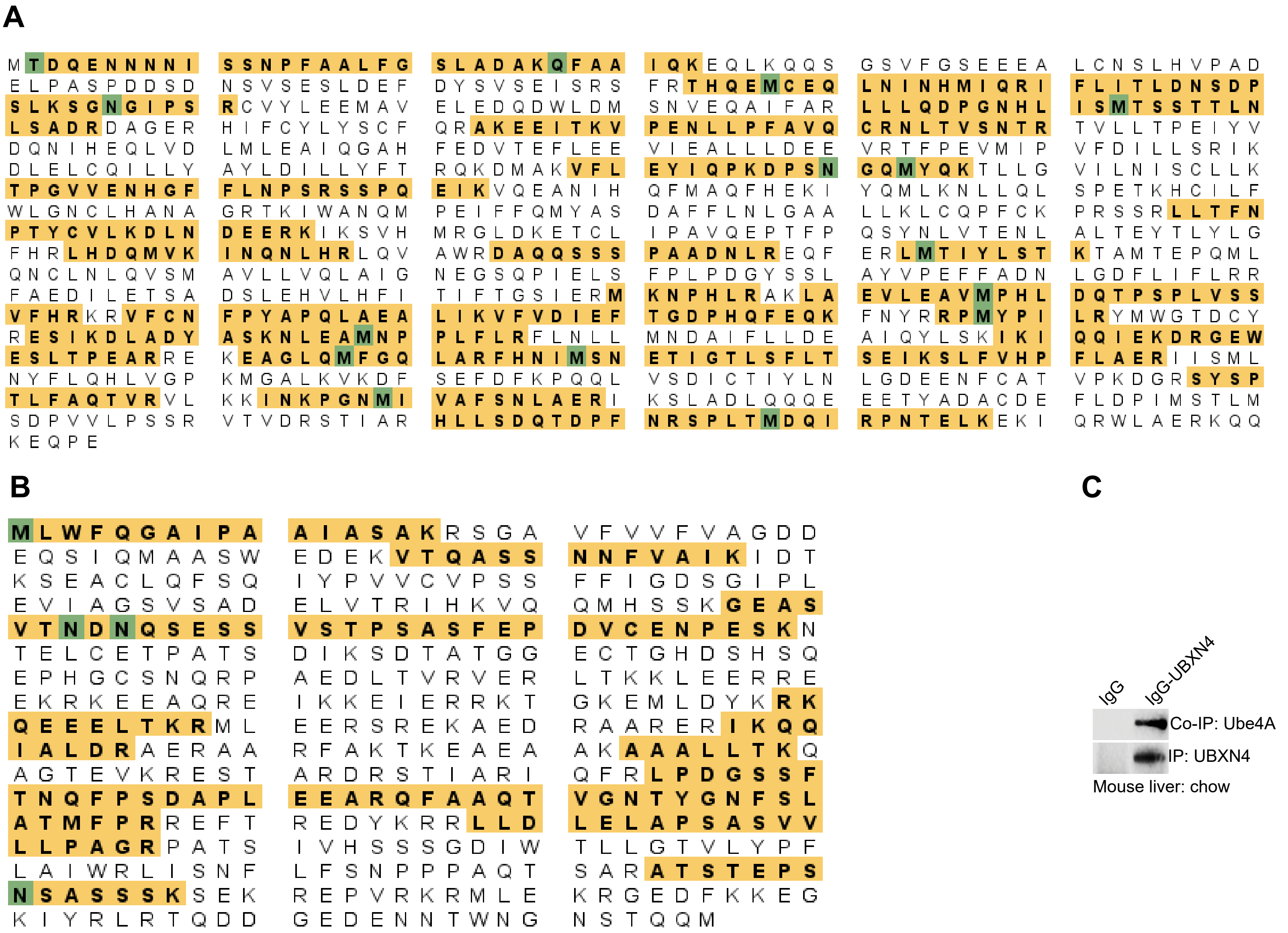


D

| Ube4A-UBXN4 interaction in HEK293 cells | |
|---|---|
| Numbers = Exclusive, unique spectrum counts | |
| Complete medium | |
| Ube4A-IP | 55 |
| UBXN4-Co-IP | 5 |

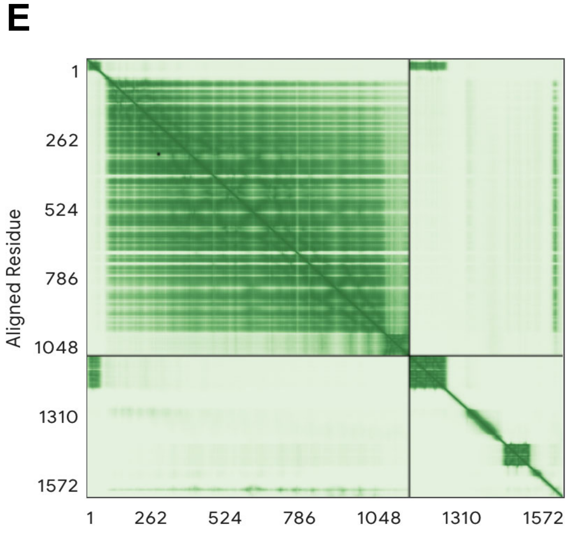


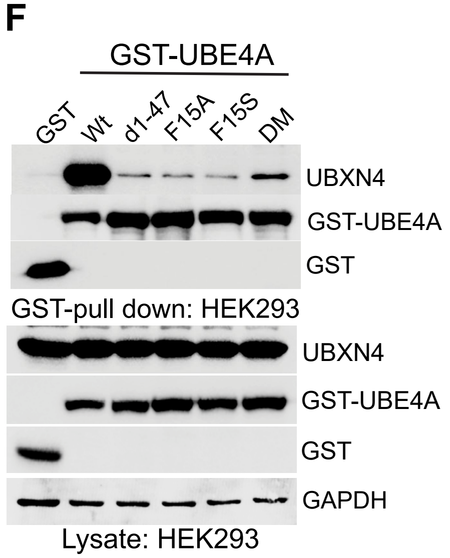
